# TDP-43 suppression of ATP8A2 cryptic splicing implicates phosphatidylserine-driven neuroinflammation in ALS/FTD

**DOI:** 10.1101/2025.11.21.689833

**Authors:** James T. O’Connor, Hui Qi Loo, Caiwei Guo, Sarah Pickles, Shea Sundali, Vidhya Maheswari Jawahar, Dennis W. Dickson, A. Joseph Bloom, Leonard Petrucelli, Aaron D. Gitler, Jeffrey Milbrandt, Aaron DiAntonio

## Abstract

Inappropriate externalization of phosphatidylserine (PS) is a candidate mechanism of pathogenic neuroinflammation, a critical driver of neurodegenerative disease. ATP8A2, a flippase that maintains PS on the plasma membrane inner leaflet, is mutated in both *Wabbler-lethal* mice and patients with the ataxia syndrome CAMRQ4. Here, we identify *ATP8A2* as a target of TDP-43 cryptic exon suppression, and demonstrate that *ATP8A2* loss leads to immune-mediated neurodegeneration. *ATP8A2* splicing is significantly dysregulated following TDP-43 depletion in human neurons and in brains of patients with Amyotrophic Lateral Sclerosis-Frontotemporal Dementia (ALS-FTD). In mice, *Atp8a2* loss increases PS exposure and promotes neuroinflammation. Depletion of peripheral macrophages rescues motor axon degeneration and doubles *Atp8a2* knockout mouse lifespan, while depletion of both peripheral macrophages and central microglia quadruples lifespan and improves coordination. Hence, ATP8A2 is a pathologically relevant TDP-43 target and inhibition of phagocytic immune cell attack against neurons is a potential treatment for patients with CAMRQ4 and ALS-FTD.

## Introduction

TDP-43 pathology—a hallmark of ALS/FTD and Limbic-predominant Age-related TDP-43 Encephalopathy (LATE) and frequent co-pathology in Alzheimer’s and Parkinson’s disease^1–8^—disrupts RNA processing, notably by promoting cryptic exon inclusion that reduces expression of key neuronal genes^9,10^. To find disease-relevant TDP-43 targets beyond classic axon/synapse genes (i.e. *STMN2*^11–15^ and *UNC13A*^16–18^), we compared RNA alterations across disparate models of TDP-43 dysfunction and found consistent mis-splicing of *ATP8A2*, a neuron-enriched^19^ phosphatidylserine (PS) flippase that maintains PS on the inner plasma membrane leaflet^20–23^ . Externalized PS is an “eat me” signal that marks unhealthy cells for engulfment by phagocytes^24–27^, a plausible driver of neuroinflammation, an important disease mechanism in TDP-43-dependent neurodegenerative diseases^28–30^. Hence, ATP8A2 presents a candidate bridge from RNA dysregulation to immune-mediated pathology.

PS exposure is sufficient to trigger dendrite fragmentation and glial phagocytosis of *Drosophila* neurons^24,31^, and loss of the pan-flippase chaperone protein CDC50 (*TMEM30A*) in Purkinje cells causes early-onset ataxia^32^. Moreover, we demonstrated that the SARM1-dependent programmed axon degeneration pathway induces PS externalization, and this enhances axon loss^33,34^. Human and animal genetics further underscore the importance of ATP8A2. Loss of ATP8A2 causes the rare neurodevelopmental disorder Cerebellar Ataxia, Impaired Intellectual Development, and Disequilibrium Syndrome Type 4 (CAMRQ4)^35–38^ whose symptoms—severe ataxia, hypotonia, and increased pediatric mortality—are recapitulated by the *Wabbler-lethal* spontaneous *Atp8a2* knockout mouse^21,39^. However, prior mammalian studies did not evaluate the role of ATP8A2 in neuroinflammation.

Here we demonstrate a dramatic increase in *ATP8A2* cryptic exon inclusion and concomitant loss of the canonical transcript upon TDP-43 knockdown in cultured human neurons and in ALS-FTD patient brains, most strikingly in progranulin (GRN+) FTD, a subtype with pronounced neuroinflammation^40–42^. Because TDP-43-regulated transcripts are species-specific^9,10^, we modeled the pathological consequences of ATP8A2 loss using *Atp8a2* knockout mice. The absence of ATP8A2 results in increased PS exposure and neuroinflammation in both the peripheral and central nervous systems. Depletion of peripheral macrophages rescues motor axon loss and doubles the lifespan of *Atp8a2* knockout mice, demonstrating that macrophages are driving the axon degeneration. Depletion of both microglia and peripheral macrophages, even after disease onset, increases lifespan more than four-fold, and improves coordination. These findings identify *ATP8A2* as a pathologically-relevant TDP-43 splicing target whose loss induces lethal neuroinflammation and neurodegeneration, and suggest that inhibition of phagocytic immune cell attack against neurons is a potential treatment strategy for patients with CAMRQ4 and ALS/FTD.

## Results

### *ATP8A2* splicing is regulated by TDP-43 in ALS-FTD

Analyzing differentially expressed RNAs in two disparate models of TDP-43 dysfunction^16,17^, we noted only 23 genes identified as both significantly downregulated in cortical-like i^3^Neurons and cryptically spliced in TDP-43-negative nuclei from ALS-FTD patient brains (Suppl. Table 1). These include Stathmin-2 (*STMN2*), *UNC13A*, and the PS flippase *ATP8A2*. Analyzing results from seven previously published ALS-FTD brains^16,43^, we found increased cryptic splicing between *ATP8A2* exons 13 and 14 compared to neurons from control brains (Figure 1A, A’). To test directly whether *ATP8A2* splicing is regulated by TDP-43, we performed RT-qPCR on human-embryonic stem cell (hESC)-derived neurons with or without TDP-43 knockdown, and found a 70% decrease in canonically-spliced transcripts and a ∼5,000-fold increase in cryptic-spliced *ATP8A2* (Figure 1B, B’), an effect enhanced to 15,000-fold by preventing nonsense-mediated decay with the hSMG-1 inhibitor 11j (Figure 1B, B’). To determine whether the *ATP8A2* transcript is similarly affected in disease, we analyzed brains from 190 FTLD patients with TDP-43 inclusions compared to 50 controls and found canonically-spliced *ATP8A2* significantly decreased and the cryptic-spliced transcript dramatically increased (Figure 1C, C’). Increased cryptic exon inclusion was present across all FTLD patient subtypes, including sporadic, C9orf72, and GRN+ (Figure 1D’). Interestingly, samples from GRN+ FTD, a subtype with prominent neuroinflammation^40–42^, showed the greatest decrease in canonical *ATP8A2* (Figure 1D). Further, canonical *ATP8A2* loss and cryptic exon inclusion were correlated with patient phosphorylated TDP-43 levels (Figure S1), a pathological hallmark of TDP-43 cytoplasmic aggregation^44^. Taken together, these data demonstrate that *ATP8A2* is a bona fide TDP-43 splicing target whose expression is disrupted in FTLD.

**Figure 1:**
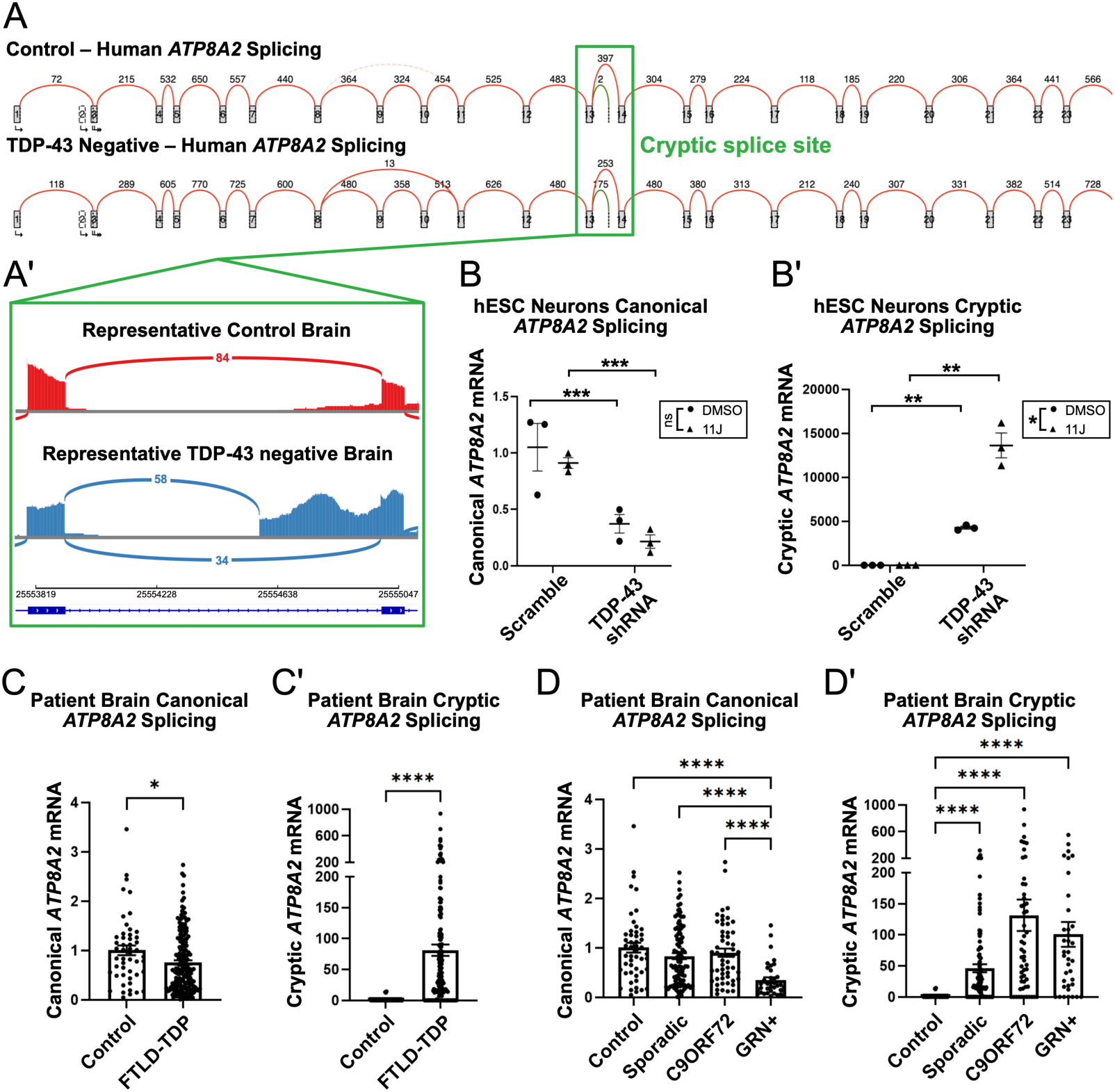
*ATP8A2* splicing is dysregulated by loss of TDP-43. A) Sashimi plot of normal splicing (red) or cryptic splicing (green) of *ATP8A2* in previously published^16,43^ neuronal nuclei of control human brains or TDP-43 negative neuronal nuclei of ALS-FTD patient brains, with the apparent cryptic splice site (green box) occurring between exons 13 and 14. Numbers represent total aggregate reads mapped to each exon. A’) Splicing between exon 13 and 14 in a representative control (red) or ALS-FTD patient (blue) brain. B) RT-qPCR data in 12-day control or TDP-43 knockdown hESC-derived neurons showing relative expression of normally spliced *ATP8A2* (B) or cryptic-spliced *ATP8A2* (B’). Enhanced expression of cryptic exons is observed after treatment with hSMG-1 Inhibitor 11j to prevent nonsense-mediated decay of cryptic transcripts. C–D) RT-qPCR data from brain cortices of 50 control patients or 190 patients who suffered from Frontotemporal Lobar Degeneration with TDP-43 Pathology (FLTD-TDP) showing relative expression of normally spliced *ATP8A2* (C-D) or cryptic-spliced *ATP8A2* (C’-D’), either combined (C) or divided by FTLD subtype (D). B) *p < 0.05, **p < 0.01, ***p < 0.001 by 2-way ANOVA C) * p < 0.05, **** p < 0.0001 by Mann-Whitney Test, D) *p < 0.05, ***p < 0.001, ****p < 0.0001 by Kruskal-Wallis Test.

### Neuroinflammation precedes motor axon degeneration in *Atp8a2* knockout mice

The *Atp8a2*-mutant *Wabbler-lethal* mouse model is characterized by a hallmark wobbling gait, a lifespan of approximately two months, and degeneration of axons in the brain and motor axons in the femoral and sciatic nerves^21,39^. Because ATP8A2 is a flippase that maintains PS on the inner leaflet of the plasma membrane^20–23^, and externalized PS is an “eat me” signal to phagocytes^24–27^, we hypothesized that this neurodegeneration is secondary to neuroinflammation triggered by aberrantly exposed PS. To assess whether ATP8A2 loss causes peripheral neuroinflammation prior to motor axon degeneration, we stained sciatic nerves of 1-month old wild-type and *Atp8a2* knockout mice for markers of neuroinflammatory macrophages and observed a significant increase in both Iba1+ macrophages and CD68+ activated macrophages, demonstrating severe peripheral nerve inflammation (Figure S2A-C). At this age, although we observe very little axon degeneration in their femoral nerves (Figure S2D-F), *Atp8a2* knockout mice already display reduced body weight (Figure S2G) and severe behavioral phenotypes including significantly decreased ability to cross a 12mm-wide catwalk (Figure S2H).

### ATP8A2 loss drives neuroinflammation and motor axon degeneration independent of SARM1

We observed severe peripheral nerve inflammation in both the sciatic (Figure 2A-B, D) and motor-predominant femoral (Figure S3A-B) nerves of *Atp8a2* knockout mice, as well as axon loss in the femoral nerve at two months of age (Figure 2E, F, H). Because ATP8A2 loss could plausibly induce axonal metabolic stress that activates the central executioner of axon degeneration, SARM1^45^, we tested the necessity for SARM1 in ATP8A2-associated degeneration utilizing a *Atp8a2*/*Sarm1* double-knockout mouse. However, neither sciatic nerve inflammation (Fig. 2C-D) nor femoral axon degeneration (Fig. 2G-H) was prevented in the *Atp8a2/Sarm1* double-knockout model, demonstrating that ATP8A2 loss induces neuroinflammation and motor axon degeneration independent of SARM1-mediated Wallerian degeneration.

**Figure 2:**
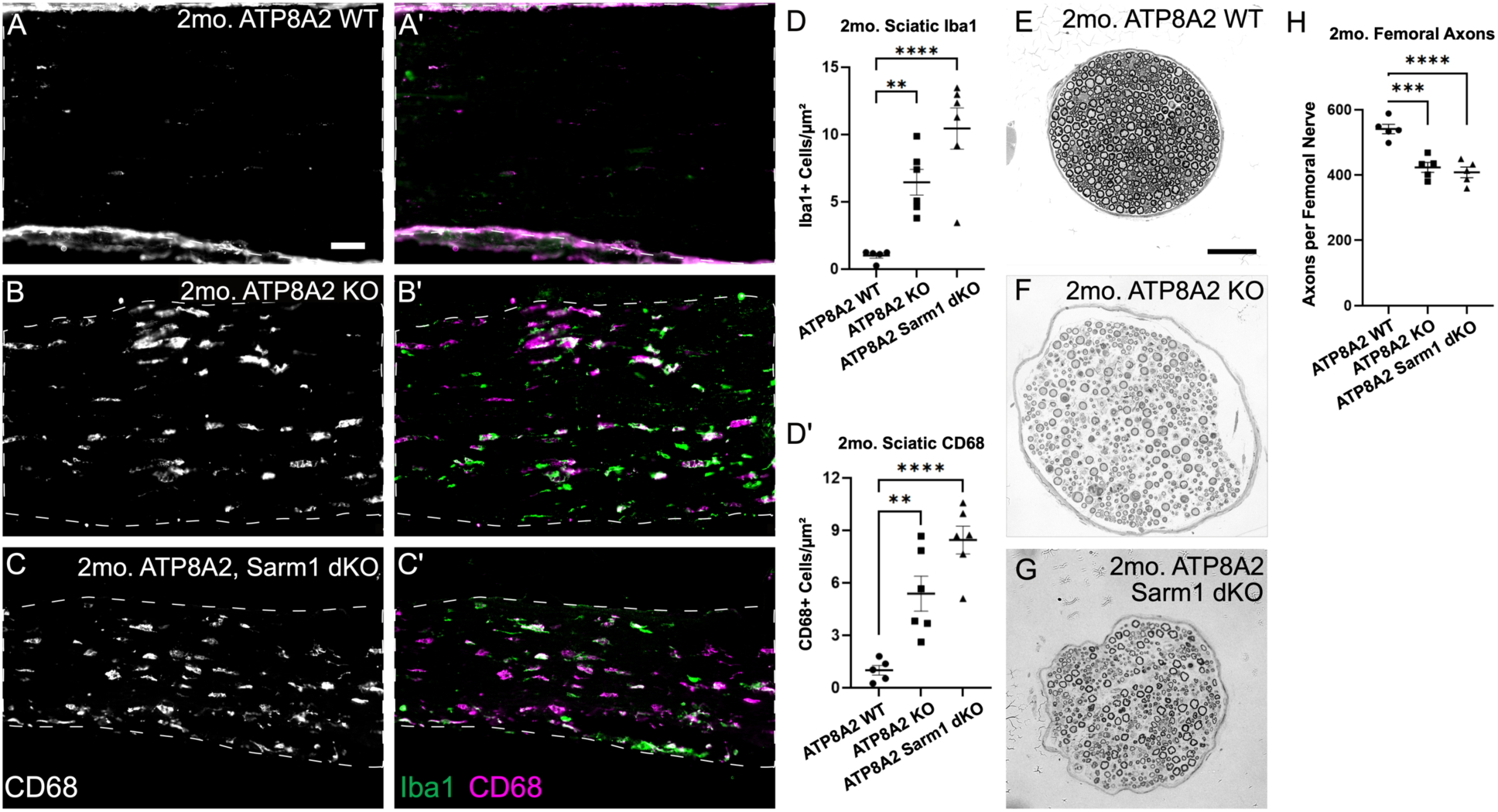
Loss of ATP8A2 induces nerve neuroinflammation and non-SARM1 dependent axon degeneration. A–D) Staining of sciatic nerves of 2-month-old mice for Iba1 (pan-macrophages, green) and CD68 (activated macrophages, magenta) shows peripheral inflammation in *Atp8a2* knockout mice (B). Knockout of *Sarm1,* to prevent the canonical cell-autonomous axon degeneration pathway, does not prevent activated macrophages from infiltrating peripheral nerves of *Atp8a2* knockout mice (C). D–D’) Quantification of Iba1+ cells (D) and CD68+ cells (D’) normalized to controls. E–H) Toluidine blue staining shows axon degeneration in *Atp8a2* knockout (F) compared to control motor-predominant femoral nerves (E), which is not rescued by knockout of *Sarm1* (G). H) Quantification of axons per nerve shows significant degeneration in two-month-old *Atp8a2* knockout femoral nerves. Scale bars = 50 µm. D, D’, H) ** p < 0.01, *** p < 0.001, **** p < 0.0001 by One-way ANOVA.

### Peripheral macrophages are necessary for motor axon loss and reduced survival in the *Atp8a2* knockout

Activated macrophages may be driving the degeneration in *Atp8a2* knockout mice or they may be simply responding to degenerating axons. To distinguish between these possibilities, we depleted macrophages with Colony Stimulating Factor 1 Receptor antibody (anti-CSF1R)^46^ on days 30 (when the mice are symptomatic but prior to femoral axon degeneration) and 45, then assessed neuroinflammation and neuropathology on day 60. The treatment successfully depleted peripheral nerve macrophages (Fig 3A-C) and, excitingly, prevented the femoral axon degeneration observed in IgG-treated controls (Fig 3D-F). Moreover, with longitudinal dosing every 15 days from day 30, median survival nearly doubled from 75 days to 148 days, with all treated knockouts outliving all controls (range 103–160 vs. 32–98 days, respectively) (Fig. 3G). Both the efficacy of macrophage depletion and axon protection also persisted at five months, with treated *Atp8a2* knockouts femoral nerves being indistinguishable from wild type (Figure S4A-F). However, despite preserving peripheral axons, the motor coordination deficits remained (Fig. 3H), implicating additional CNS drivers.

**Figure 3:**
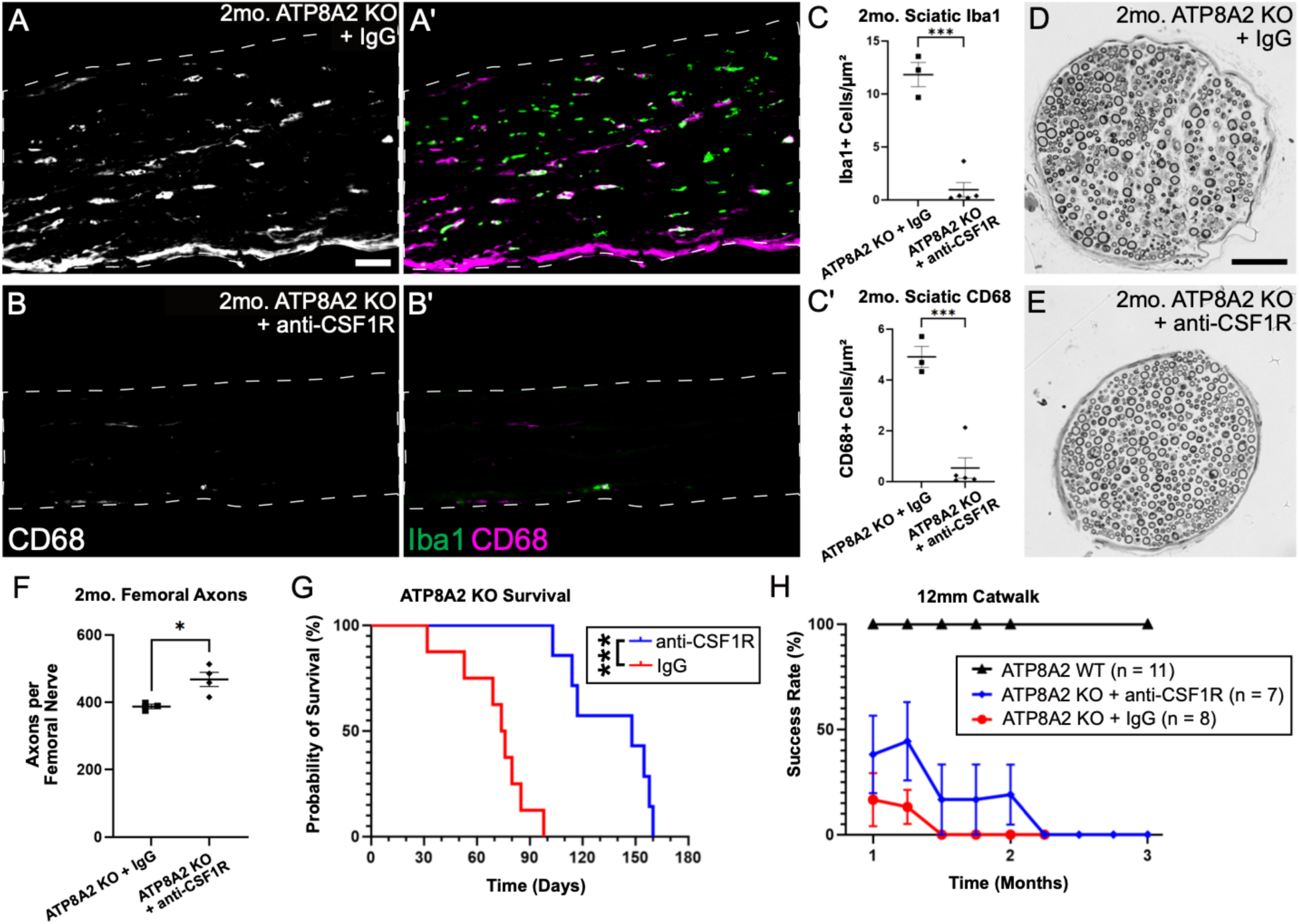
Peripheral macrophage depletion via CSF1R antibody rescues motor axon degeneration and doubles lifespan of *Atp8a2* knockout mice. A–C) Staining of sciatic nerves of 2-month mice for Iba1 (pan-macrophages, green) and CD68 (activated macrophages, magenta) shows peripheral macrophage depletion in *Atp8a2* knockout mice treated with anti-CSF1R at Day 30 and 45 (B) compared to IgG control (A). C) Quantification of Iba1+ cells (C) and CD68+ cells (C’). D–F) Toluidine blue staining shows axon degeneration in *Atp8a2* knockout motor-predominant femoral nerves treated with IgG (D) that is rescued after treatment with anti-CSF1R (E). F) Quantification of axons per nerve shows significant prevention of axon degeneration after treatment with anti-CSF1R. G) Survival curve of *Atp8a2* knockout mice treated every 15 days, beginning at Day 30, showing an improvement in lifespan from a median survival time of 75 days (min: 32, max: 98) with IgG (red), to 148 days (min: 103, max: 160) with anti-CSF1R (blue). H) 12-millimeter catwalk test shows no improvement in mice treated with IgG or anti-CSF1R from one month of age. Scale bar = 50 µm. C, C’, F) * p < 0.01, *** p < 0.001 by Student’s t-test. G) *** p < 0.001 by Mantel-Cox Test.

### Brain-penetrant CSF1R inhibition dramatically prolongs the lifespan of *Atp8a2* knockout mice

To evaluate the degree to which ATP8A2 loss causes neuroinflammation in the brain, we performed flow cytometry from brains of control and 1-month-old *Atp8a2* knockout mice. We observed a ∼20-fold expansion of microglia expressing high levels of the damage-associated marker CLEC7A and low levels of the homeostatic marker P2RY12, both hallmarks of disease-associated microglia (DAMs)^47–49^ (Figure 4A-B). We further stained sagittal brain sections for Iba1 and CD68, markers of phagocytic microglia, focusing on regions near the deep cerebellar nuclei–a pathological locus in *Wabbler-lethal* mice^21^–and the cerebellar peduncles, containing efferent cerebellar axons associated with balance and coordination^50^ (Figure S5A, red box). Iba1+ microglia and CD68+ microglial lysosomes are significantly increased in the cerebellar peduncles of *Atp8a2* knockout mice (Figure 4C–D, Figure S5B–C), indicating the presence of activated, phagocytic microglia.

**Figure 4:**
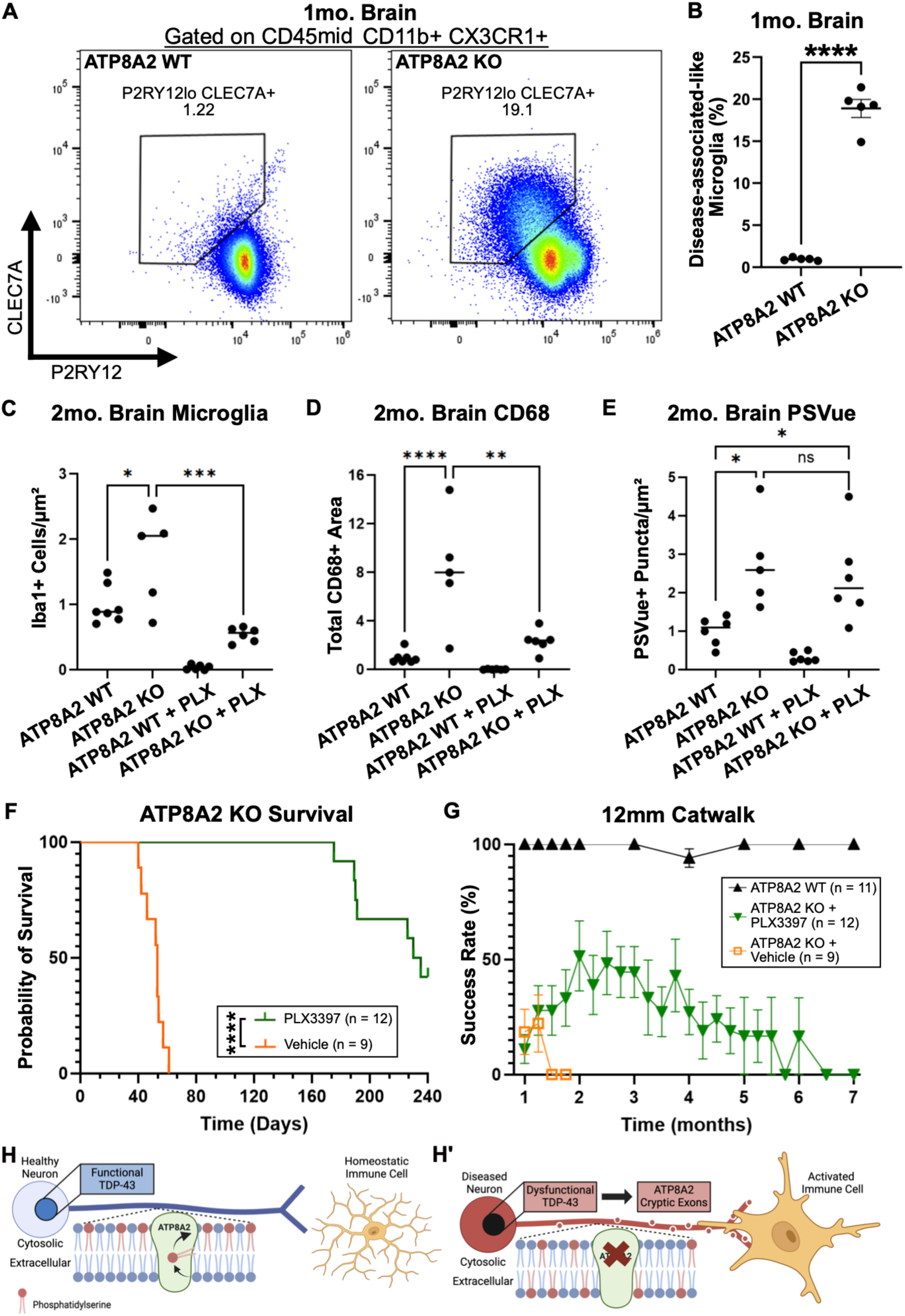
Depletion of disease-associated microglia quadruples survival and improves behavior of *Atp8a2* knockout mice. A) Representative flow cytometry plots of 1-month brains of control (left) or *Atp8a2* knockout (right) mice gated for CD45-mid, CD11b+, CX3CR1+ microglia, comparing levels of the damage-associated marker CLEC7A and the homeostatic marker P2RY12. B) Quantification of CLEC7A+ P2RY12-microglia showing a significant increase in disease-associated microglia in 1-month *Atp8a2* knockout brains. n = 5 for each genotype. C–E) Quantification (corresponding to images in Figures S5 and S6) of Iba1+ cells per area (C), CD68+ puncta area (D), or PSVue puncta per area (E) in the cerebellar peduncles of 2-month control or *Atp8a2* knockouts, treated with or without 600ppm PLX3397, showing loss of ATP8A2 significantly increases microglia (C), neuroinflammation (D), and externalized PS (E). PLX3397 treatment is sufficient to significantly decrease inflammation but leaves externalized PS unchanged in *Atp8a2* knockout mice. F) Survival curve of PLX3397-treated (n = 6 at 300ppm, 6 at 600ppm) *Atp8a2* knockout mice with a median survival time of 235 days compared to vehicle fed mice (n = 9) with a median survival time of 53 days. G) 12-millimeter catwalk test shows vehicle mice unable to complete this task after 45 days of age, while PLX-fed mice show behavioral improvement in the first month of treatment, which gradually tapers off over time. H) Hypothesized model of a healthy (H) or diseased (H’) neuron whereby disease-induced neuroinflammation is caused by loss of functional ATP8A2 after TDP-43 loss, leading to aberrantly externalized phosphatidylserine and improper axonal phagocytosis. B) **** p < 0.0001 by Student’s t-test. C–E) * p < 0.05, ** p < 0.01, *** p < 0.001, **** p < 0.0001 by One-way ANOVA. F) **** p < 0.0001 by Mantel-Cox Test.

To determine whether CNS inflammation contributes to the *Atp8a2* knockout phenotypes, we treated 30-day old mice with the brain-penetrant CSF1R inhibitor PLX3397, which depletes microglia^51,52^. 30-days-old *Atp8a2* knockout mice already show significant neurological defects, and so we wished to treat younger mice, but beginning treatment at 21-days-old caused seizures and rapid death. Instead, we treated with 600ppm PLX3397, a dose previously used to rapidly deplete microglia^52^, beginning at day 30 and observed a significant decrease in the number of microglia and CD68+ puncta around the cerebellar peduncles in *Atp8a2* knockout mice (Figure 4C–D, Figure S5D–E). Interestingly, *Atp8a2* knockouts appeared more resistant to microglial depletion than controls (∼70% reduction vs 96% reduction), a phenotype previously observed in other severe neuroinflammatory models^53^. To assess whether loss of ATP8A2 leads to accumulation of externalized PS as expected from its molecular function, we stained wild type and knockout brains. Before fixation, brains were submerged in PSVue-643, which binds to externalized phosphatidylserine, and observed a significant increase of PSVue puncta in the cerebellar peduncles of *Atp8a2* knockout brains—an increase unaffected by PLX3397 treatment (Figure 4E, Figure S6)—confirming that loss of ATP8A2 increases externalized PS. Finally, we observed that treatment with PLX3397 depletes peripheral macrophages from sciatic nerves of 2-month *Atp8a2* knockout mice (Figure S7), demonstrating its sufficiency to target both peripheral and central neuroinflammation.

Mice treated with 300 or 600 ppm PLX3397 were combined into one cohort because they achieved comparable improvements in lifespan and coordination, with no side effects other than expected loss of fur pigmentation^53^. PLX3397 treatment from day 30 dramatically extended the median lifespan of *Atp8a2* knockout mice from 53 days (range: 42–60) to 235 days (range: 175-240+) and significantly improved narrow catwalk performance (Fig. 4F–G). Together, these findings demonstrate that *Atp8a2* knockout behavioral phenotypes are driven by both central and peripheral neuroinflammation that can be ameliorated by neuroimmune modulators even after disease onset and support a model in which ATP8A2 loss—whether congenital or TDP-43-associated—leads to aberrant phosphatidylserine exposure provoking a massive immune cell attack on the nervous system (Figure 4H).

## Discussion

TDP-43 proteinopathy is the defining lesion of ALS and LATE, and common in FTLD and Alzheimer’s Disease^1–9^. This broad association with neurodegenerative disease underscores the centrality of TDP-43 to nervous system homeostasis. Cytoplasmic TDP-43 drives gain-of-function toxicity—mitochondrial dysfunction, aberrant stress granule formation, reduced autophagy, and decreased proteasome function^5,9^—while loss of nuclear TDP-43 disrupts RNA-processing^54,55^, which is thought to contribute to disease through various gene-specific mechanisms ^9,10^. This is exemplified by the mis-splicing of transcripts for the axonal and synaptic proteins *STMN2*^11,14^*, UNC13A*^16,17^, and the potassium channel *KCNQ2*^56^. Loss of the endogenous transcripts for these genes disrupts the axonal cytoskeleton, NMJ maintenance, axon regeneration, synaptic function, and neuronal excitability, all potential contributors to ALS pathology^13,15,18,56–58^. Here we nominate the flippase ATP8A2, a neuronal regulator of the phagocytic ‘eat-me’ signal phosphatidylserine^21,31^, as a driver of the prominent neuroinflammatory component of TDP-43 pathology^28,29^. We identify a reproducible cryptic exon event in *ATP8A2* after acute TDP-43 knockdown in iPSC-derived neurons and, critically, in ALS-FTD brains. Neuroinflammation is best studied *in vivo*, but TDP-43 splicing targets are species specific, so to investigate whether ATP8A2 loss induces neuroinflammation, we turned to the ATP8A2 knockout mouse. We found that ATP8A2 loss is sufficient to increase exposed PS in the brain and induce lethal neuroinflammation and motor axon degeneration; *Atp8a2* knockout mice display early macrophage infiltration followed by motor-predominant axon loss. Peripheral macrophage depletion prevents femoral motor axon degeneration and extends lifespan, while brain-penetrant CSF1R inhibition further prolongs life, improves coordination, and reduces cerebellar microgliosis. These results support a model in which TDP-43-associated ATP8A2 loss drives pathological exposure of PS, mobilizing microglial/macrophage phagocytosis, culminating in central and peripheral axon degeneration and premature death.

Our findings have immediate clinical implications for CAMRQ4, a rare syndrome resulting from mutations in ATP8A2, with symptoms including severe intellectual disability, ataxia, blindness, and pediatric death^35,36,59^. Importantly, the CSF1R inhibitor PLX3397 was found to be safe and generally well-tolerated in phase I and II human clinical trials against glioblastoma^60,61^. Our results provide evidence that immunomodulation, by PLX3397 or another agent, could be an effective therapy for CAMRQ4 patients. Further therapeutic development could improve upon broad immunosuppression by more precisely targeting dysregulated phosphatidylserine externalization at two levels: 1) the immune cells, by targeting receptors that respond to PS and 2) the exposing cells, by neutralizing or cloaking the externalized PS. Importantly, while CAMRQ4 is rare, neurodegeneration with TDP-43 involvement is common, and TDP-43-driven ATP8A2 loss could play a central role in the neuroinflammation observed in these diseases. Thus, therapies developed for CAMRQ4 could also help in ALS/FTD and other TDP-43 proteinopathies, especially in subtypes where neuroinflammation is prominent.

Although all FTD subtypes showed significantly increased *ATP8A2* cryptic splicing, by far the largest decrease in properly-spliced *ATP8A2* was observed in progranulin (GRN+) FTD. This is notable because GRN mutations account for nearly one third of non-sporadic FTD worldwide^62^, and GRN-FTD is a distinctly neuroinflammatory form characterized by gliosis, microglial activation, and neuron loss^40,41^. Although other ALS-FTD subtypes, such as C9orf72, have pro-inflammatory signatures^30,63^ correlated with disease progression^64^, GRN-FTD is particularly informative mechanistically. GRN loss in microglia increases proinflammatory cytokines and promotes neuronal death^65^, while neuronal GRN loss impairs peripheral nerve regeneration^66^, underscoring the gene’s importance across neuronal and non-neuronal cell types. Notably, progranulin-negative microglia alone are sufficient to trigger TDP-43 aggregation in neurons via C1q/C3-mediated neuroimmune crosstalk^42^. The selective loss of canonical *ATP8A2* in GRN-FTD therefore suggests a feed-forward loop whereby inflammatory microglia induce TDP-43 proteinopathy, suppressing ATP8A2 expression, which further amplifies neuroinflammation.

TDP-43 targets are not conserved in mice^9,10^, so the impact of TDP-43 pathology has been studied *in vivo* by knocking out the mouse orthologs of human TDP-43 targets, such as UNC13A^67,68^ and STMN2^12,13,15^. Because *UNC13A* and *STMN2* are commonly mis-spliced in TDP-43 proteinopathies^11,14,16,17^, they are leading therapeutic targets for antisense-oligonucleotides (ASOs) to block cryptic splicing^69^, but their loss has not been linked to neuroinflammation. Meanwhile, although ATP8A2 has not been studied in relation to common disease, the profound TDP-43-dependent *ATP8A2* mis-splicing we observe makes it a compelling target to mitigate both neuroinflammation and axon loss. While anti-inflammatory strategies have so far been largely unsuccessful in clinical trials^28,70^ despite promising mouse model data^71^, targeting a key driver of inflammation like ATP8A2 with ASOs, or via a PS-specific immunomodulator, could yield better results.

Prior work reported that the Wallerian Degeneration Slow (*Wld^S^*) mutation prevented motor axon degeneration in *Wabbler-lethal* mice, although it had no impact on behavior or survival^21^. Since *Wld^S^* suppresses SARM1 activation^45,72,73^, we were surprised that *Atp8a2/Sarm1* double-knockout mice still exhibit motor axon degeneration. There are a few potential explanations for this discrepancy. First, we used an engineered *Atp8a2* allele predicted to delete a large exon and cause a frameshift, while the previous study^21^ characterized the spontaneous *Wabbler-lethal* allele, a small in-frame deletion. However, the described phenotypes are very similar, making this explanation unlikely. Alternatively, in addition to suppressing SARM1 activation^45,72,73^, *Wld^S^* also boosts NAD^+^ synthesis^73–75^, which may protect axons from degenerating through an additional mechanism. Finally, unaccounted-for differences in mouse genetic background or husbandry may contribute. Regardless, neither *Wld^S^* nor *Sarm1* knockout improved behavior nor survival in *Atp8a2* deficient mice; only immune cell depletion achieved this, showing the neuropathy caused by ATP8A2 loss is driven primarily by neuroinflammation.

Treatment with PLX3397 beginning at 30 days of age dramatically improved outcomes in *Atp8a2* knockout mice, but neurological defects persisted. While non-immunological consequences of ATP8A2 loss could contribute, we attribute the incomplete rescue to timing; earlier depletion was not tolerated, and by 30 days significant neuroinflammatory damage had already accrued. Indeed, without any nutritional intervention, the knockout mice die at 26-32 days^21^. Targeting CSF1R genetically using the Csf1r^ΔFIRE^ allele^76^ could be a tractable alternative; by depleting phagocytic macrophages and microglia genetically, Csf1r^ΔFIRE^ may prevent early neuroinflammatory damage that occurs *in utero* and during development. A limitation of this study is that we do not yet know which cell types expose PS to trigger lethal neuroinflammation, though we favor neurons as the key player. The pattern of degeneration in CAMRQ4—peripheral motor axons^21^, the visual and auditory systems^20^, and specific brain regions—suggests a specific neuronal rather than glial subtype as the primary driver, though these cellular contributors are not mutually exclusive. Regardless, we have shown that neuroimmune attack is responsible for the disease symptoms in *Atp8a2* knockout mice, and that macrophage/microglial depletion is sufficient to block this pathology, extend lifespan, and improve behavior. Thus, identifying *ATP8A2* as a bona fide TDP-43 target opens therapeutic avenues to treat not only CAMRQ4, but ALS-FTD and other TDP-43-proteinopathies in which neuroinflammation is a central pathological mechanism.

## Acknowledgements

We thank Yo Sasaki (Washington University, St. Louis, MO) and Adriana Norris (Emory University, Atlanta, GA) for their input on this manuscript. We thank Amy Strickland, Cassidy Menendez, and Sylvia Johnson for technical support relating to this project. We thank the Mayo Clinic Brain Bank and all the deceased patients and their families for access brain tissue used in this study.

## Key resources table

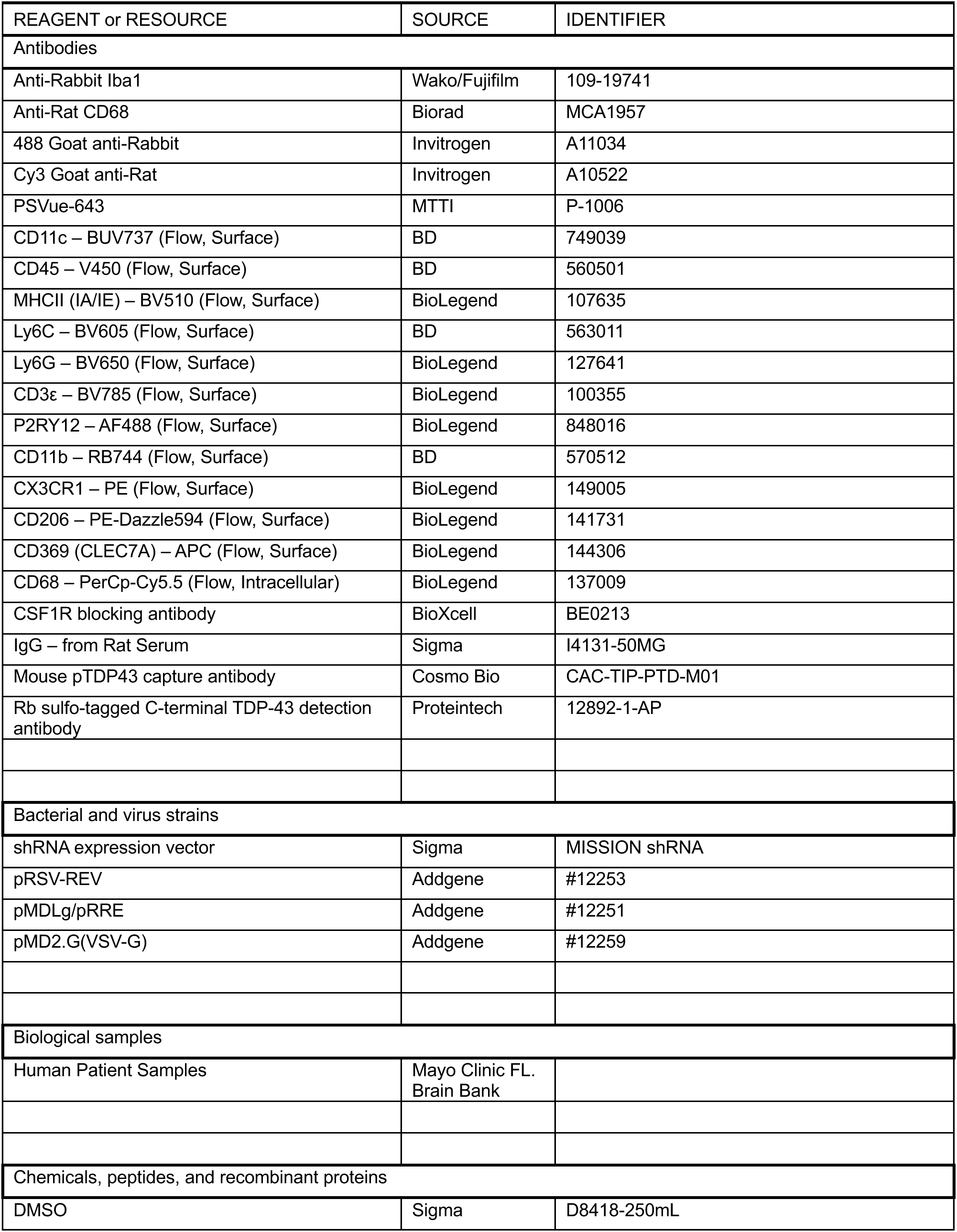
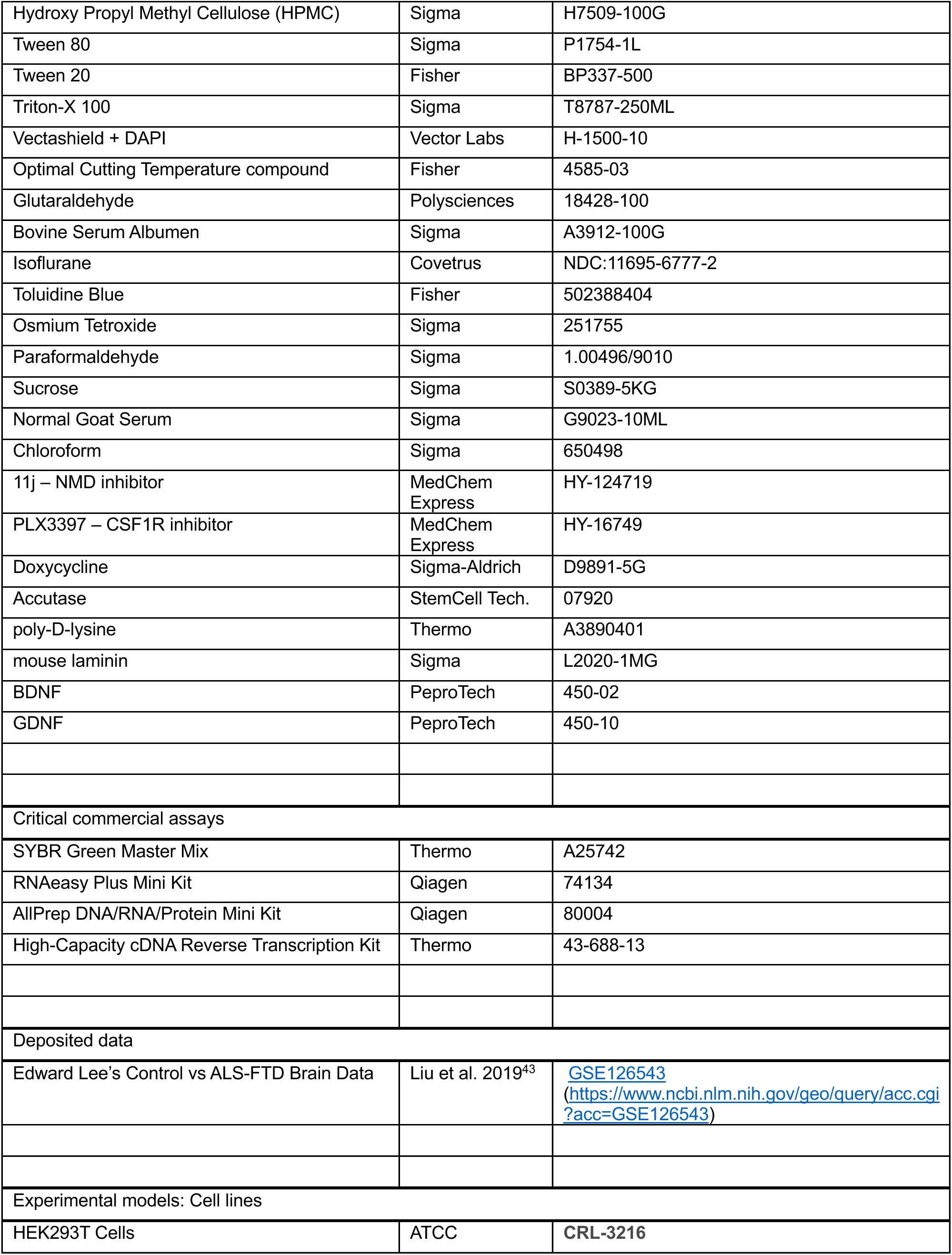
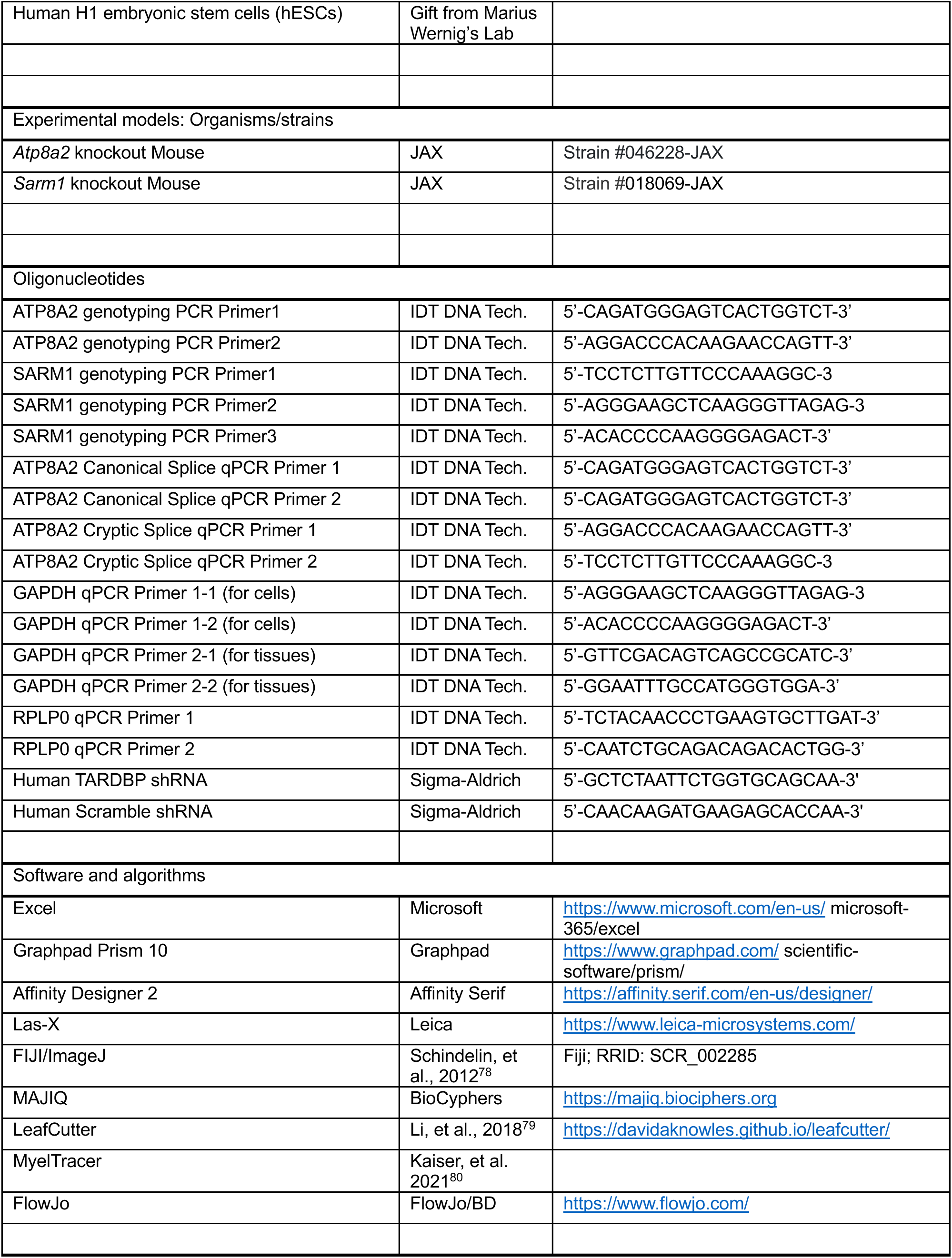
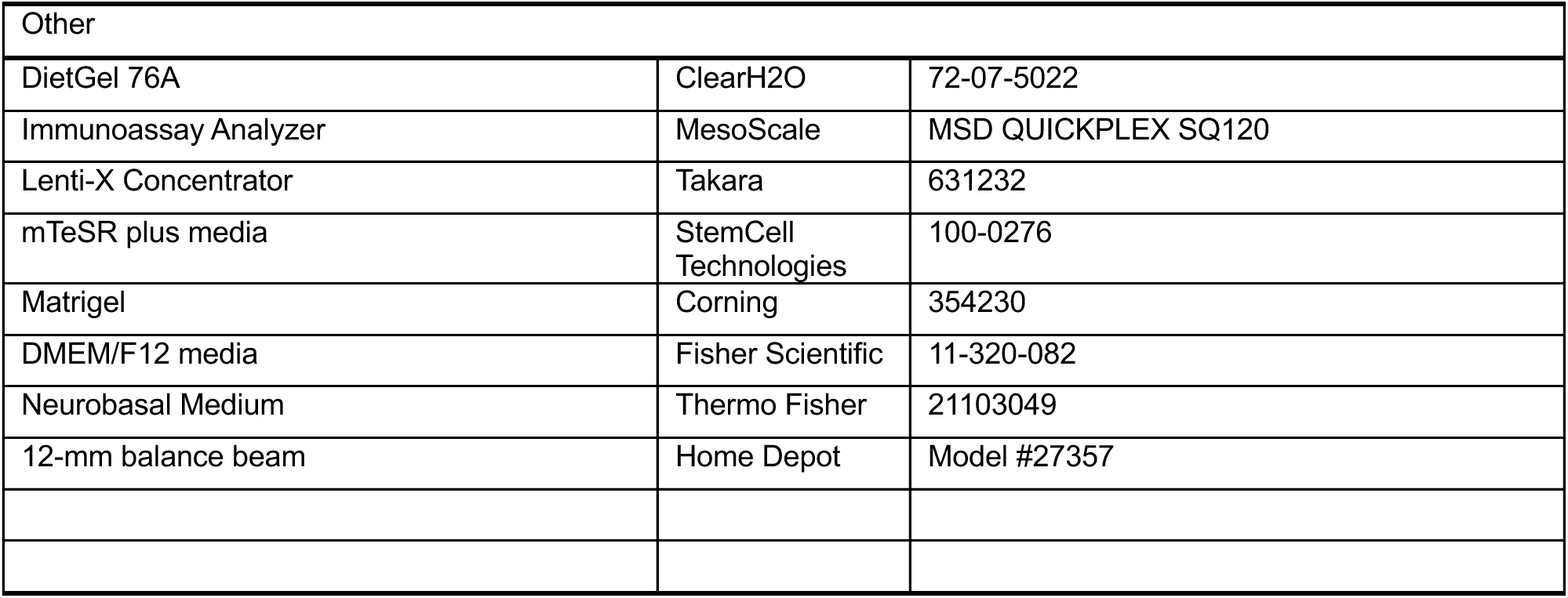

## RESOURCE AVAILABILITY

### Lead contact

Further information and requests for resources and reagents should be directed to and will be fulfilled by the lead contact, Aaron DiAntonio or co-corresponding author Jeffrey Milbrandt.

### Materials availability

No new mouse or cell lines were generated in this manuscript. All mouse lines are available from JAX.

### Data and code availability

RNA-seq data for splicing analysis in Figure 1A is available at https://www.ncbi.nlm.nih.gov/geo/query/acc.cgi?acc=GSE126543, (GSE126543)

## EXPERIMENTAL MODEL AND SUBJECT DETAILS

### Mouselines

The “*Atp8a2* heterozygous” mouse harboring a 365bp deletion in exon 2 of *Atp8a2*, C57BL/6NJ-Atp8a2em1(IMPC)J/Mmjax (JAX, MMRRC Strain #046228-JAX) was obtained from JAX. *Atp8a2* heterozygotes were bred together, and genotyped using primers:

> 5’-CAG ATG GGA GTC ACT GGT CT -3’ and

> 5’-AGG ACC CAC AAG AAC CAG TT -3’.

SARM1 knockout mice (SARM1 B6.129X1-Sarm1tm1Aidi/J The Jackson Laboratory JAX:018069, were initially received as a gift from Macro Colonna at Department of Pathology and Immunology, Washington University School of Medicine in St. Louis^81^, and were genotyped using a 3-primer system of:

> 5’-TCC TCT TGT TCC CAA AGG C -3’

> 5’-AGG GAA GCT CAA GGG TTA GAG -3 and

> 5’-ACA CCA CCA AGG GGA GAC T -3’

### Mouse Husbandry

Mice were housed and used following the institutional animal study guidelines and protocols approved by the Institutional Animal Care and Use Committee of Washington University in St. Louis. Mice were maintained on a 12-hour light-dark cycle and food/water provided ad libitum. Both female and male mice were used for the experiments described in this study. Cages containing *Atp8a2* knockout mice were given one cup/68 grams (or 1.5 cups/102 grams if mice consistently finished their food before next feeding) of DietGel76A (ClearH2O, SKU: 72-07-5022) every two days on the floor of their cage.

## METHOD DETAILS

### Macrophage Depletion

IgG (from rat serum, reagent grade, ≥95% (SDS-PAGE), essentially salt-free, lyophilized powder, Millipore Sigma I4131-50MG) was dissolved in pure water to match the anti-Csf1R concentration of 9.5mg/mL. 1.5 mg (∼160µL) of anti-CSF1R (InVivoMab anti-mouse CSF1R (CD115), BioXcell BE0213), or IgG was injected intraperitoneally into each mouse at 15-day intervals beginning at Day 30.

### Microglia Depletion

600ppm PLX3397 in mouse chow was previously used to rapidly and effectively deplete microglia^52^. When fed via oral gavage, 65mg/kg/day of PLX5622 (approximately the same molecular weight as PLX3397) was considered nominally close to a 300ppm dose of PLX in chow, to effectively deplete microglia^82,83^. Therefore, to feed five 12-gram mice for two days with PLX3397 mixed in DietGel76A required ∼7.5mg of drug for each 300ppm cup of food or ∼15mg of drug for each 600ppm cup of food.

PLX3397 (Pexidartinib, MedChemExpress HY-16749) was dissolved in DMSO at 0.167g/mL and stored in small aliquots at -20°C. To add PLX3397 into DietGel76A required careful mixing of ingredients in order from hydrophobic to hydrophilic to prevent the drug from precipitating out of solution. For a 300ppm cup of PLX food, 240µL of Tween80 (Sigma, P1754-1L) was mixed first with 40mg of hydroxypropyl methylcellulose (Sigma, H7509-100G). 45µL of PLX3397 in DMSO (Sigma, D8414-250mL) at 0.167g/mL was then added to the Tween80/HPMC mixture, which was thoroughly mixed via vortex. Finally, 720µL of water was added and vigorously vortexed to mix the drug into an amphipathic solution that did not crash out. For a 600ppm dose, this recipe was doubled. The PLX mixture was then added to 1.5 cups (102 grams) of DietGel76A that had been melted in a microwave for ∼30 seconds, mixed thoroughly, then allowed to cool. For vehicle mice, this process was repeated using pure DMSO rather than PLX3397 in DMSO.

### Mouse Harvesting

Mice were anesthetized with isoflurane (Covetrus, NDC 11695-6777-2) and then perfused with 10-20mL of ice-cold PBS through the heart until the lungs and liver appeared clear. The brain was taken, bisected along the corpus collosum, and both hemispheres were placed into 4% paraformaldehyde (Sigma, 1.00596.9010) at 4°C for 24 hours, and then transferred to sucrose for multiple days for cryopreservation. Both sets of sciatic, sural, femoral, and tibial nerves were removed and placed on toothpicks with a marking to indicate proximal and distal side of the nerve and placed into 4% paraformaldehyde at 4°C for at least 1 hour. Nerves used for immunostaining were transferred to 30% sucrose (Sigma S0289-5KG) for multiple days for cryopreservation followed by freezing with Fisher TissuePlus OCT (Fisher 4585-03) into cryomold blocks (Sakura TissueTek 15x15x5mm, Ref 4566) for cryosectioning, while nerves used for plastic sectioning were transferred to 3% glutaraldehyde (diluted from Polysciences 18428-100) for further fixation. Mouse tails were taken to re-test and confirm each mouse’s genotype.

### Nerve Immunofluorescence staining

Nerves were cryosectioned on a Leica cryostat (Leica CM1950) with a thickness of 6µm on Superfrost Plus slides. Nerves were permeabilized with ice-cold acetone for 10 minutes, washed in PBS, and blocked for 1 hour at room temperature with PBS + 0.1% Triton-X100 (Sigma T8787-250ML) + 4% BSA (Sigma A3912-100G). Primary antibodies Iba1 (anti-rabbit 1:500, WAKO/Fujifilm 109-19741), CD68 (anti-rat 1:100, Biorad MCA1957) were diluted in block and placed on slides overnight at 4°C. The following day, slides were washed multiple times with PBS + 0.1% Triton-X, then secondary antibodies 488 goat-anti rabbit (Invitrogen A11034) or Cy3 goat-anti rat (Invitrogen A10522) were added to PBS + 0.1% Triton-X for one hour at room temperature. The slides were then washed multiple times with PBS + 0.1% Triton-X, before adding vectashield+DAPI (Vector Labs H-1500-10) and covering with coverslips (Corning 24x50mm-1thickness, Cat# 2975-245).

### Brain Immunofluorescence staining

Brains were cryosectioned on a Leica cryostat (Leica CM1950) with a thickness of 30µm on Superfrost Plus slides using sagittal cuts. Nerves were permeabilized with ice-cold acetone for 10 minutes, washed in PBS + 0.1% Tween20 (Fisher BP337-500), and blocked for 1.5 hours at room temperature with PBS + 0.1% Tween20+ 5% Normal Goat Serum. Primary antibodies Iba1 (anti-rabbit 1:500, WAKO/Fujifilm 109-19741), CD68 (anti-rat 1:100, Biorad MCA1957) were diluted in block and placed on slides overnight at room temperature. The following day, slides were washed multiple times with PBS + 0.1% Tween20, then secondary antibodies 488 goat-anti rabbit (Invitrogen A11034) or Cy3 goat-anti rat (Invitrogen A10522) were added to PBS + 0.1% Tween20 for two hours at room temperature. The slides were then washed multiple times with PBS + 0.1% Tween20, before adding vectashield+DAPI (Vector Labs H-1500-10) and covering with coverslips (Corning 24x50mm-1thickness, Cat# 2975-245).

### Brain PSVue staining

Immediately upon harvesting, one hemisphere of mouse brain was placed into an eppendorf tube containing 10 µM PSVue-643 (MTTI, Cat#P-1006) in PBS for 20 minutes while shaking. The brains were then washed with PBS before fixation and sucrose dehydration as normal, with the brains continuously protected from light. PSVue signal was found to remain even after cryosectioning and antibody staining.

### Plastic sectioning and toluidine blue staining

Briefly, nerves were fixed in 3% glutaraldehyde in 0.1 mL PBS overnight at 4C, washed and stained with 1% osmium tetroxide (Sigma 251755) overnight at 4C. Nerves were washed and dehydrated in a serial gradient of 50% to 100% ethanol. Nerves were then incubated in 50% propylene oxide/50% ethanol, then 100% propylene oxide. Following that, nerves were incubated in Araldite resin/propylene oxide solutions in 50:50, 70:30, 90:10 ratios for 24 h, and subsequently embedded in 100% Araldite resin solution (Araldite: DDSA: DMP30, 12:9:1, Electron Microscopy Sciences) and baked at 60C overnight. Semithin 400–600 nm sections were cut using a Leica Ultramicrotome (Leica EM UC7), placed on microscopy slides, and stained with Toluidine blue (Fisher 502388404). Staining and quantification were performed using the Myeltracer.

### Cell isolation and flow cytometry analysis

Mice were euthanized and transcardially perfused with 20mL ice-cold PBS. Half hemispheres of the brains were dissected out and kept in ice-cold PBS until homogenization. Brain tissues were cut into smaller pieces with a razor blade and then transferred to a 7mL Dounce tissue grinder (DWK Life Sciences, Cat# 8853000007) with 7mL ice-cold FACS buffer (PBS + 0.5% BSA). The tissues were homogenized by 20 strokes with pestle A followed 20 strokes with pestle B. The homogenate is then filtered through a 70mm filter (Fisher, Cat# 50-233-5745) into 50mL tubes pre-coated with FACS buffer. The tubes were centrifuged at 400 x g for 5 mins, the supernatant discarded, and the pellet resuspended with 10mL of 30% isotonic Percoll (Cytiva, Cat# 17-0891-02) in 15mL tubes. The tubes were centrifuged at 800 x g (acceleration 6, brake 1) for 15 mins. The myelin layer and supernatant were removed by vacuum suction, and the cell pellets were resuspended in 5mL FACS buffer. The tubes were centrifuged at 350 x g for 5 mins, the supernatant discarded, and the cell pellets resuspended in 200mL PBS and transferred to a 96-well U-bottom plate. The plate was centrifuged at 500 x g for 5 mins and the supernatant discarded. The cells were resuspended in 100mL PBS containing Fixable Viability 780 (Invitrogen Cat# 65-0865-14) diluted at 1:5000 for 20 mins. The cells were washed in FACS buffer and centrifuged at 500 x g for 5 mins. The cells were resuspended in 50mL FACS buffer containing anti-CD16/32 (BioLegend, Cat# 101302) diluted 1:50 for 5 mins. The cells were then stained by adding another 50mL of an antibody cocktail to make up final concentrations of 20% Brilliant Stain Buffer Plus (BD Biosciences, Cat# 566385) in FACS buffer and the cell surface antibodies listed in Supplemental Table 1 for 30 mins on ice. The cells were washed twice in FACS buffer and fixed with 4% PFA on ice for 20 mins. The cells were washed twice in FACS buffer and stored at 4°C overnight. On the second day, the cells were resuspended in 1X Perm/Wash buffer (BD Biosciences, Cat# 554714) for 15 mins at on ice. The plate was centrifuged at 500 x g for 5 mins and the supernatant discarded. The cells were then stained with intracellular antibodies in 1X Perm/Wash buffer (Supplemental Table 1) for 30 mins on ice. The cells were washed twice in 1X Perm/Wash buffer. The cells are finally resuspended in FACS buffer and transferred to 5mL polystyrene tubes (Fisher, Cat# 14-959-2A). The samples were analyzed on a 5-laser Cytek Aurora spectral analyzer. Results were analyzed with with FlowJo v10 (BD Biosciences).

### Microscopy

Mouse tissues were imaged on a Leica Thunder Imager (Leica Microsystems, Inc.) running LAS-X software. Immunofluorescence images of nerves were taken on a 20x/0.6NA objective. Ultrathin sections of femoral axons were taken on a 40x/0.95NA objective. Brain images were taken on a 40x/0.95NA objective after an initial scan in the DAPI channel using a 5x/0.12NA objective was used to identify the region of interest.

### Catwalk Test

The catwalk test was modified from Luong, et al. 2011^84^. Briefly, a 12 mm x 4 cm x 91 cm poplar wooden board (a yard-stick) (Home Depot, Model #27357) was used as a balance beam, suspended over a cushioned basket, with one end resting on a biosafety hood. Marks were made at ∼30 cm intervals to divide the beam into thirds. Mice were placed on the first mark facing towards the biosafety hood, mice were given one minute to walk the ∼60 cm across the beam to the hood. Failure to complete this task, either by clinging to the bar, failing to reach the end, or falling off entirely, was considered a failure, while reaching the end within 60 seconds was considered a success. Mice were picked up by the tail and turned around if they attempted to go backwards, and gently prodded forwards if they paused on the beam.

### Stem cell culture

Human H1 embryonic stem cells (hESCs) were maintained in mTeSR plus media (StemCell Technologies, 100-0276) on plates coated with Matrigel (Corning, 354230), and passaged every 3-4 days using ReLeSR (StemCell Technologies, 100-0483). hESCs were differentiated into cortical-like neurons (iNeurons) by overexpressing transcription factor neurogenin 2 (NGN2) driven by a Tet-On induction system in DMEM/F12 media supplemented with 1 ug/mL doxycycline, as previously described (Bieri et al., 2019). Cells were dissociated using Accutase (StemCell Technologies, 07920) on day 3 of differentiation and replated on plates coated with poly-D-lysine (Thermo Scientific, A3890401) and mouse laminin (Sigma-Aldrich, L2020-1MG) in Neurobasal Medium (Thermo Fisher, 21103049) containing neurotrophic factors, BDNF and GDNF (R&D Systems). After four days of seeding, iNeurons were infected with shRNA lentivirus.

### shRNA and lentiviral packaging

Lentiviral packaging was achieved by transfecting HEK293T cells with each shRNA expression vector (Sigma-Aldrich, MISSION shRNA) and third generation packaging vectors pRSV-REV, pMDLg/pRRE, pMD2.G(VSV-G) (Addgene), followed by supernatant collection and concentration using Lenti-X Concentrator (Takara, 631232). Viral titers were determined by serial dilution followed by puromycin viability test. shRNA sequence for TARDBP is GCTCTAATTCTGGTGCAGCAA and for scramble control is CAACAAGATGAAGAGCACCAA.

### Inhibition of the nonsense-mediated decay

After treating scramble shRNA or TDP-43 shRNA virus for seven days, iNeurons and iMNs were treated with 0.5uM of 11j (MedChem Express, HY-124719), a SMG-1 inhibitor, or an equal volume of DMSO for 24 hours, as previously described^85^.

### RNA extraction and RT-qPCR (cell culture)

Total RNA from iNeurons was extracted using AllPrep DNA/RNA/Protein Mini Kit (Qiagen, 80004) per manufacturer’s instructions. Reverse transcription into cDNA was conducted by High-Capacity cDNA Reverse Transcription Kit (Thermo Fisher Scientific, 43-688-13). RT-qPCR was performed using Applied Biosystems PowerUp SYBR Green Master Mix (Thermo Fisher Scientific, A25742) on a QuantStudio 3 Real-Time PCR System (Thermo Fisher Scientific). The primers are:

> GAPDH-F: GAAGGTGAAGGTCGGAGTC,

> GAPDH-R: GAAGATGGTGATGGGATTTC,

> ATP8A2-cryptic-F: ACAACAATCTTATTCCCATCAGTCT,

> ATP8A2-cryptic-R: CAAGAGATGGAAGTTGCAGTGA,

> ATP8A2-canonical-F: TTCATAAACTGGGACACAGATA,

> ATP8A2-canonical-R: CGTTCCAGTCTTGTCAGAAA.

### Human post-mortem tissues

All human brain tissue was provided by the Mayo Clinic Florida Brain Bank. Written informed consent from all individuals or their family members was obtained prior to collection and all protocols associated with tissue collections were approved by the Mayo Clinic Institutional Review Board and Ethic Committee. Both cognitively normal individuals and those with a diagnosis of Frontotemporal lobar degeneration with TDP-43 pathology, as evaluated by trained neurologists and neuropathologists, were included in this study. The overall sample size was determined by availability of tissues and quality of RNA extracted from those tissues. Information regarding patient characteristics can be found in Table S1.

### RNA extraction, cDNA synthesis and qRT-PCR (human tissues)

RNA was extracted from frozen human postmortem frontal cortex using the RNAeasy Plus Mini Kit (QIAGEN). RNA integrity number (RIN) was assessed using an Agilent 2100 bioanalyzer (Agilent Technologies), and only RNA samples with a RIN > than 7.0 or higher went on to cDNA synthesis. cDNA was synthesized from 1000 ng of RNA using the High-Capacity cDNA Reverse Transcription Kit (Applied Biosystems). For quantitative reverse transcription-PCR (qRT-PCR), cDNA samples were run in triplicate, using SYBR GreenER qPCR SuperMix (Invitrogen) and run on a Prism 7900HT Fast Real-Time system (Applied Biosystems). Results were analyzed using a modified ΔΔCt method, normalized to the endogenous controls (*RPLP0* and *GAPDH*), and then normalized to levels of controls. Primers sequences are noted below:

> *RPLP0* forward: 5’-TCTACAACCCTGAAGTGCTTGAT-3’

> *RPLP0* reverse: 5’-CAATCTGCAGACAGACACTGG-3’

> *GAPDH* forward: 5’-GTTCGACAGTCAGCCGCATC-3’

> *GAPDH* reverse: 5’-GGAATTTGCCATGGGTGGA-3’

> *ATP8A2* canonical forward: 5’-TTCATAAACTGGGACACAGATA-3’

> *ATP8A2* canonical reverse: 5’-CGTTCCAGTCTTGTCAGAAA-3’

> *ATP8A2* cryptic forward: 5’-ACAACAATCTTATTCCCATCAGTCT-3’

> *ATP8A2* cryptic reverse: 5’-CAAGAGATGGAAGTTGCAGTGA-3’

### Immunoassays for phosphorylated TDP-43

Phosphorylated TDP-43 protein levels in the urea-soluble fraction from postmortem human frontal cortex tissue from individuals with FTLD-TDP and controls were previously measured by sandwich Meso Scale Discovery (MSD) immunoassay^77^ using a capture and detection antibody for phosphorylated TDP-43. Briefly, samples were diluted in Tris buffered saline (TBS) in duplicate and response values corresponding to the intensity of emitted light upon electrochemical stimulation of the assay plate using the MSD QUICKPLEX SQ120 were acquired.

## QUANTIFICATION AND STATISTICAL ANALYSIS

### Peripheral Macrophage Quantification

Iba1+ and CD68+ Macrophages in nerves were counted by hand using the Cell Counter plugin in FIJI and divided by the total area of the nerve measured using the DAPI signal. The values of cells/area were then normalized by the average control value of each group. Data were analyzed via Student’s T-test (if 2 groups were compared) or One-Way ANOVA (if >2 groups were compared) in Graphpad Prism.

### Femoral Axon Quantification

Femoral axons were analyzed by Myeltracer in a blinded manner for axon count and axon diameter size distribution. Data were analyzed via Student’s T-test (if 2 groups were compared) or One-Way ANOVA (if >2 groups were compared) in Graphpad Prism.

### Brain Inflammation Quantification

Iba1+ microglia and PSVue+ puncta in brains were counted by hand in a blinded manner using the Cell Counter plugin in FIJI and divided by the total area of the cerebellar peduncles measured using the DAPI signal. CD68+ lysosomes signal in brains were thresholded in a blinded manner, and then the total area was measured using the Measure function to determine the total area of CD68+ coverage, allowing for us to account for both plausible brain neuroinflammation phenotypes of increased lysosome number and increased size, and divided by the total area of the cerebellar peduncles measured using the DAPI signal. These values were then normalized by the average control value of each group. Data were analyzed via One-Way ANOVA in Graphpad Prism.

### RNA-seq and splicing analysis

RNA-seq data from human patient brains sorted based on the level of nuclear TDP-43 ^43^ was analyzed using MAJIQ and LeafCutter as previously described^16^.

### Figure Design

Figures were designed in Affinity Designer (Affinity Serif), with the graphical model in Figure 4H designed using BioRender.

---

**Figure S1:**
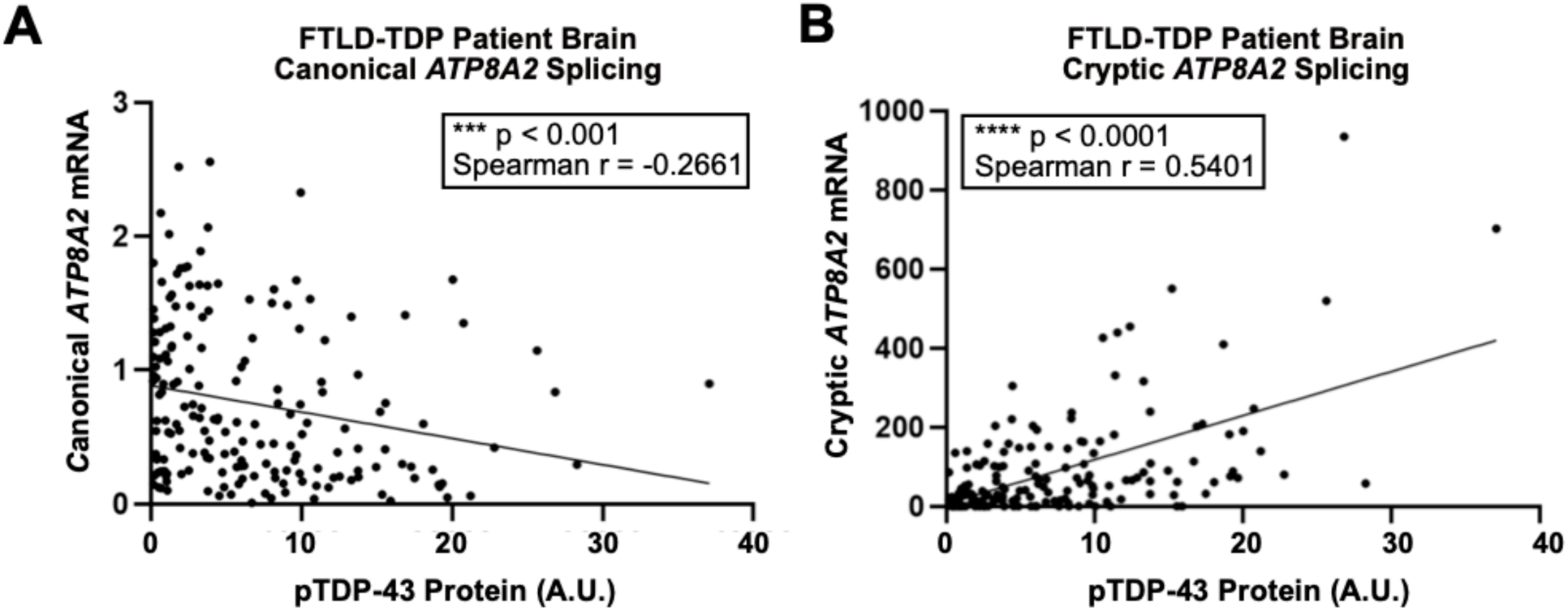
pTDP-43 Protein is correlated with dysregulation in *ATP8A2* splicing in FTLD-TDP Patient Brains. A-B) Correlation between pTDP-43 protein, measured using a Meso Scale Discovery (MSD) immunoassay^77^ and canonically-spliced *ATP8A2* mRNA (A) or cryptic-spliced *ATP8A2* mRNA (B) in 190 FTLD patient brains with TDP-43 Pathology. Canonically-spliced *ATP8A2* RNA is significantly negatively correlated (Spearman r = -0.2661, p < 0.001), while cryptic-spliced *ATP8A2* RNA is significantly positively correlated with pTDP-43 levels (Spearman r = 0.5401, p < 0.0001).

**Figure S2:**
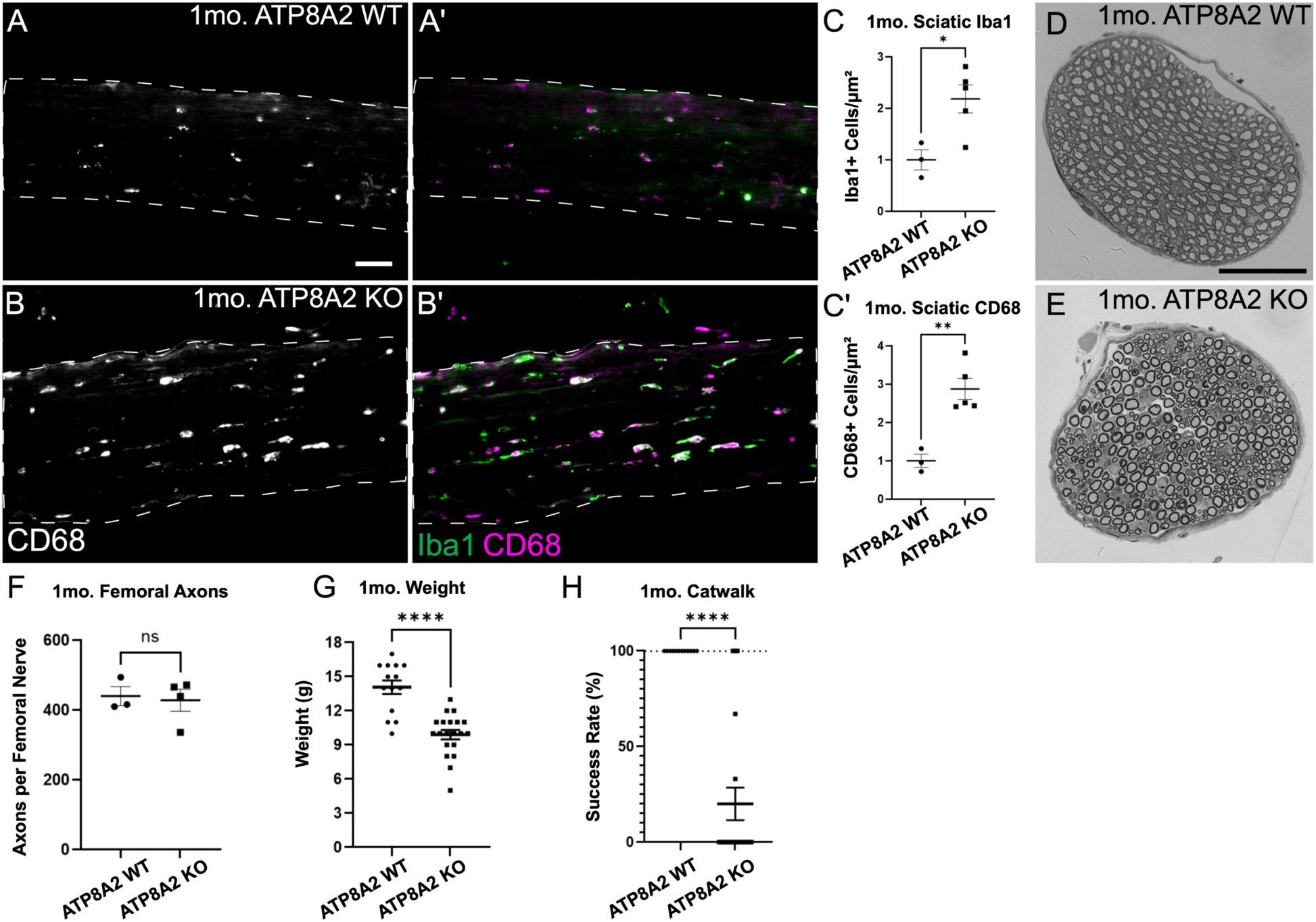
Loss of *Atp8a2* induces early neuroinflammation and behavioral defects. A–C) Staining of sciatic nerves for Iba1 (pan-macrophages, green) and CD68 (activated macrophages, magenta) shows peripheral inflammation in 1-month-old *Atp8a2* knockout (B) compared to control mice (A). C–C’) Quantification of Iba1+ cells (C) and CD68+ cells (C’) normalized to controls. D–E) Toluidine blue staining shows no significant axon degeneration in 1-month-old *Atp8a2* knockout femoral nerves (F) compared to control (E). H) Quantification of axons per nerve shows no significant degeneration in *Atp8a2* knockout femoral nerves by one month of age. G–H) One-month-old *Atp8a2* knockout mice show weight loss (G) and discoordination by catwalk test (H). Scale bars = 50 µm. C, C’, F–H) * p < 0.05, ** p < 0.01, **** p < 0.0001 by Student’s t-test.

**Figure S3:**
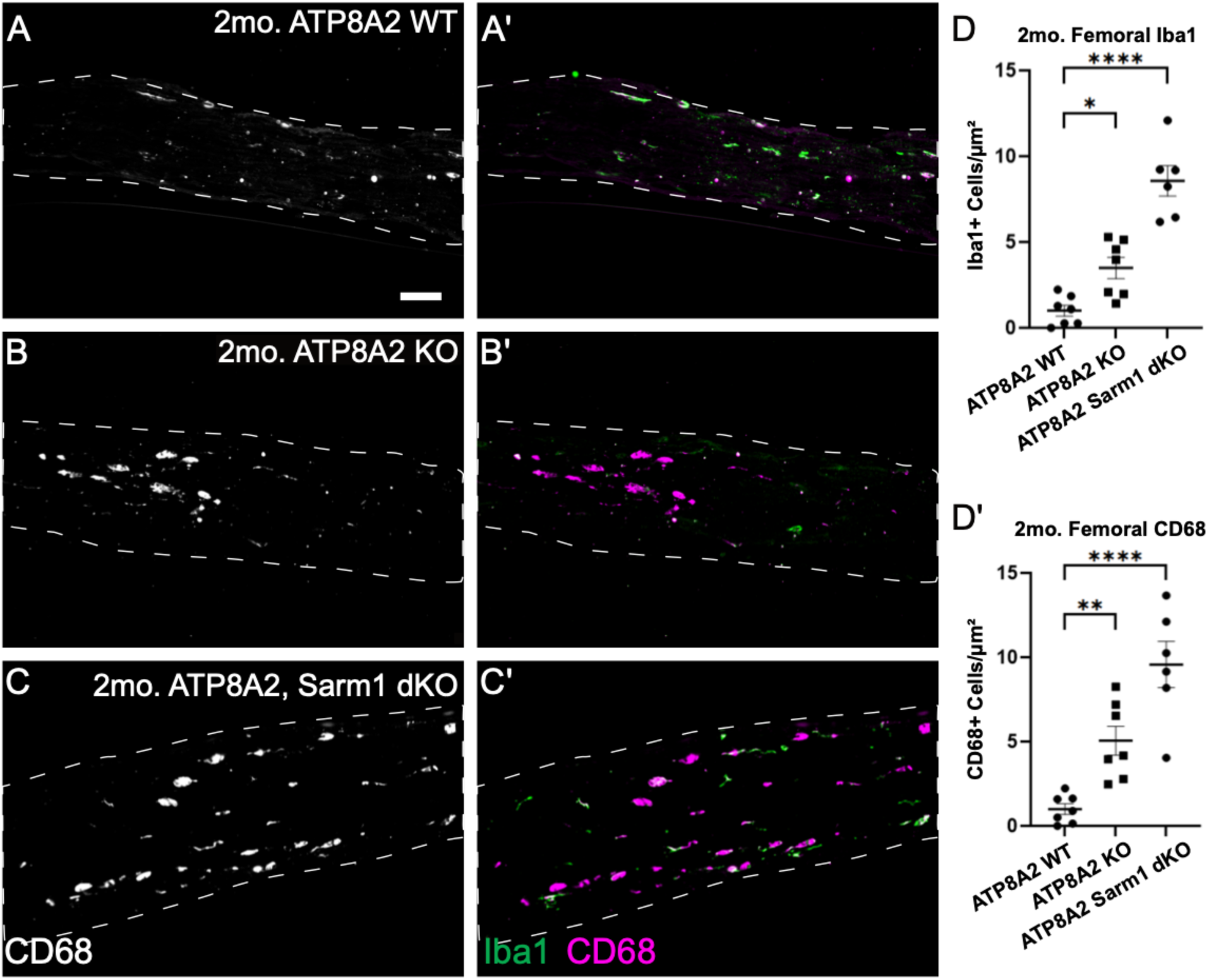
Loss of *Atp8a2* causes Sarm1-independent femoral nerve inflammation. A–D) Staining of femoral nerves of 2-month-old mice for Iba1 (pan-macrophages, green) and CD68 (activated macrophages, magenta) shows peripheral inflammation in *Atp8a2* knockout mice (B) (A). Knockout of *Sarm1* does not prevent activated macrophage infiltration of peripheral nerves (C). D–D’) Quantification of Iba1+ cells (D) and CD68+ cells (D’) normalized to controls. Scale bar = 50 µm. D, D’) * p < 0.05 ** p < 0.01, **** p < 0.0001 by One-way ANOVA.

**Figure S4:**
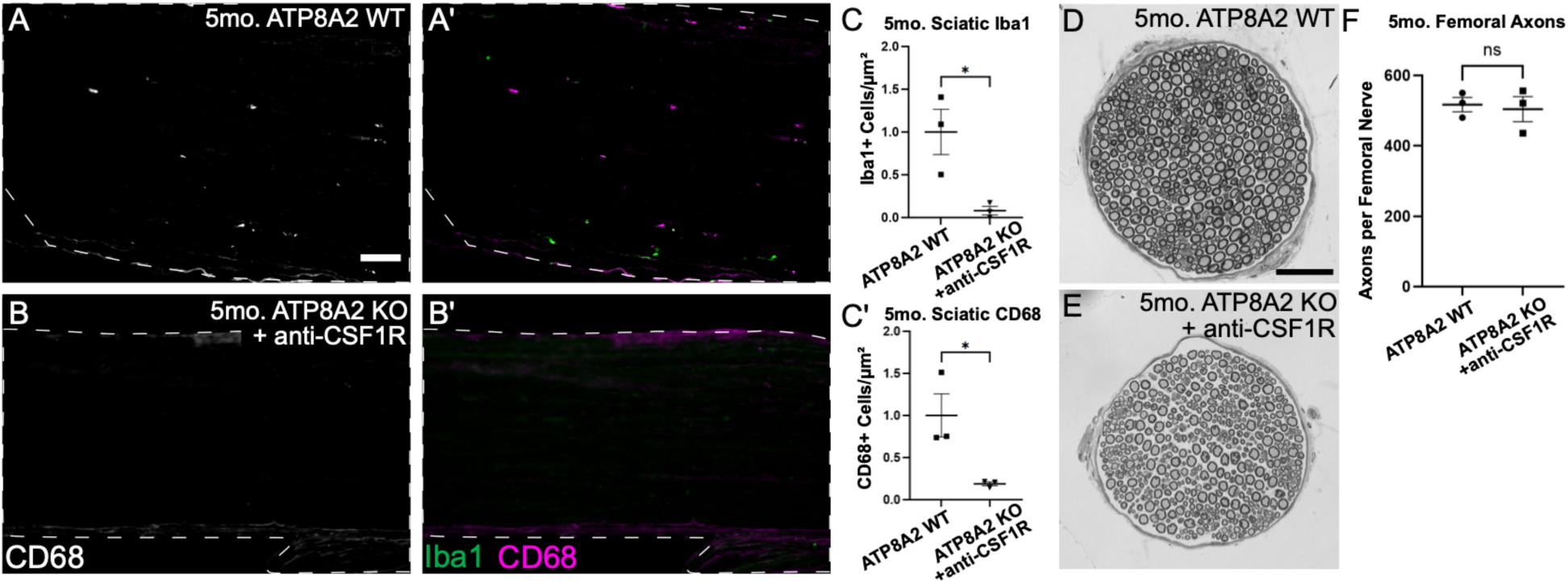
CSF1R Antibody Dosing Sustains Peripheral Macrophage Depletion for Months. A–C) Staining of Sciatic nerves of 5-month mice for Iba1 (pan-macrophages, green) and CD68 (activated macrophages, magenta) shows peripheral macrophage depletion in *Atp8a2* knockout mice treated with anti-CSF1R every 15 days beginning at Day 30, (B) even compared to control mice (A). C) Quantification of Iba1+ cells (C) and CD68+ cells (C’). D–F) Semi-thin sections of motor-predominant femoral nerves stained for toluidine blue show no significant axon degeneration in *Atp8a2* knockout femoral nerves treated with anti-CSF1R (E) compared to controls (D). F) Quantification of axons per nerve shows no significant axon degeneration in 5-month *Atp8a2* knockout nerves after continued treatment with anti-CSF1R. Scale bars = 50 µm. C, C’, F) * p < 0.05 by Student’s t-test.

**Figure S5:**
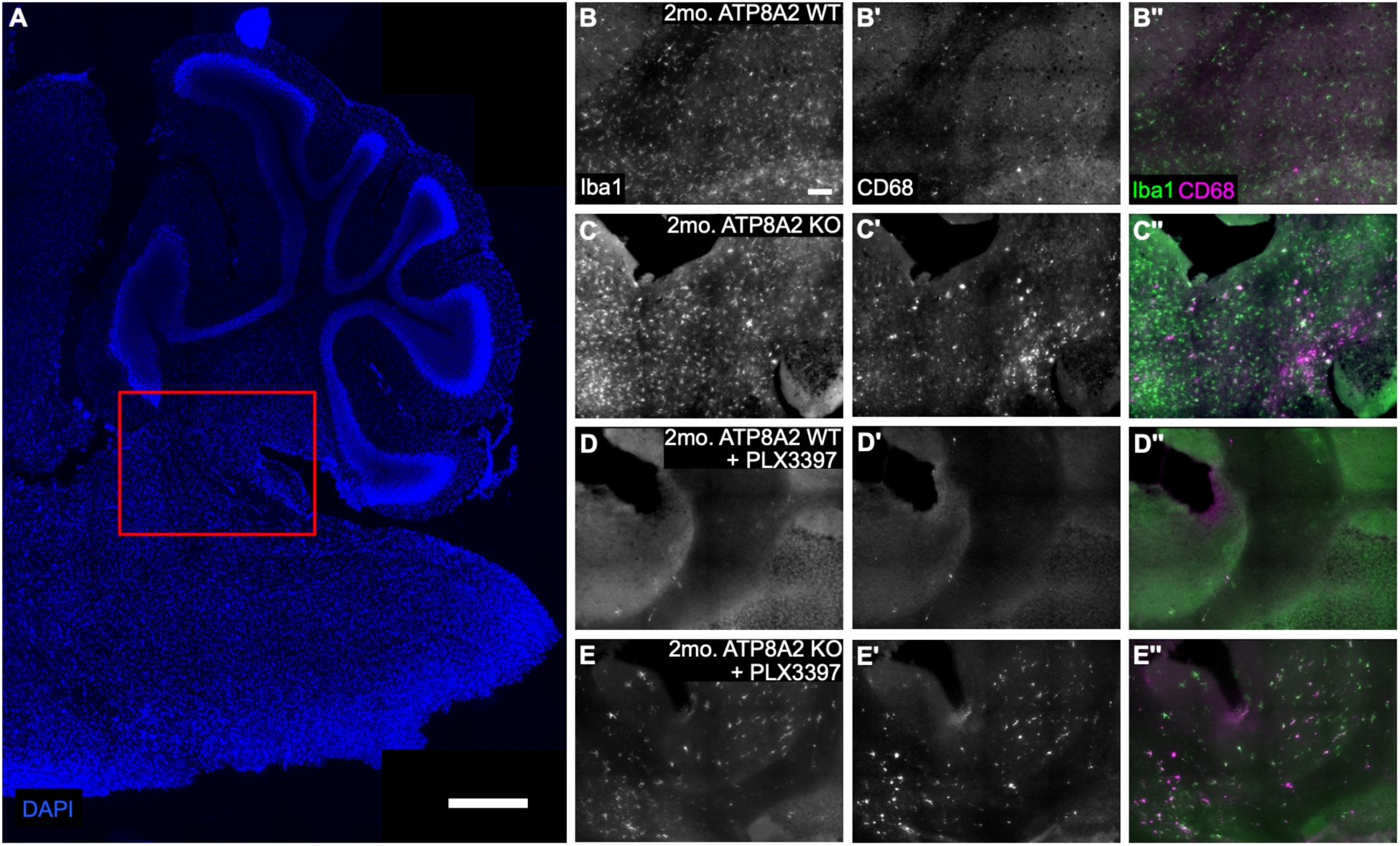
Loss of *Atp8a2* causes CNS inflammation near the cerebellar peduncles that can be rescued by PLX3397. A) Sagittal section of mouse brains stained for DAPI, showing the cerebellum, cerebellar peduncles (red box), and brainstem. Scale bar = 500 µm B-E) Brains stained for Iba1 (green) and CD68 (magenta) showing an increase in CD68 specifically in the region of the cerebellar peduncles in 2-month *Atp8a2* knockout mice compared to controls that is significantly decreased after treatment with 600ppm PLX3397. Scale bar = 50 µm in B–E. Quantification in Figure 4C, D.

**Figure S6:**
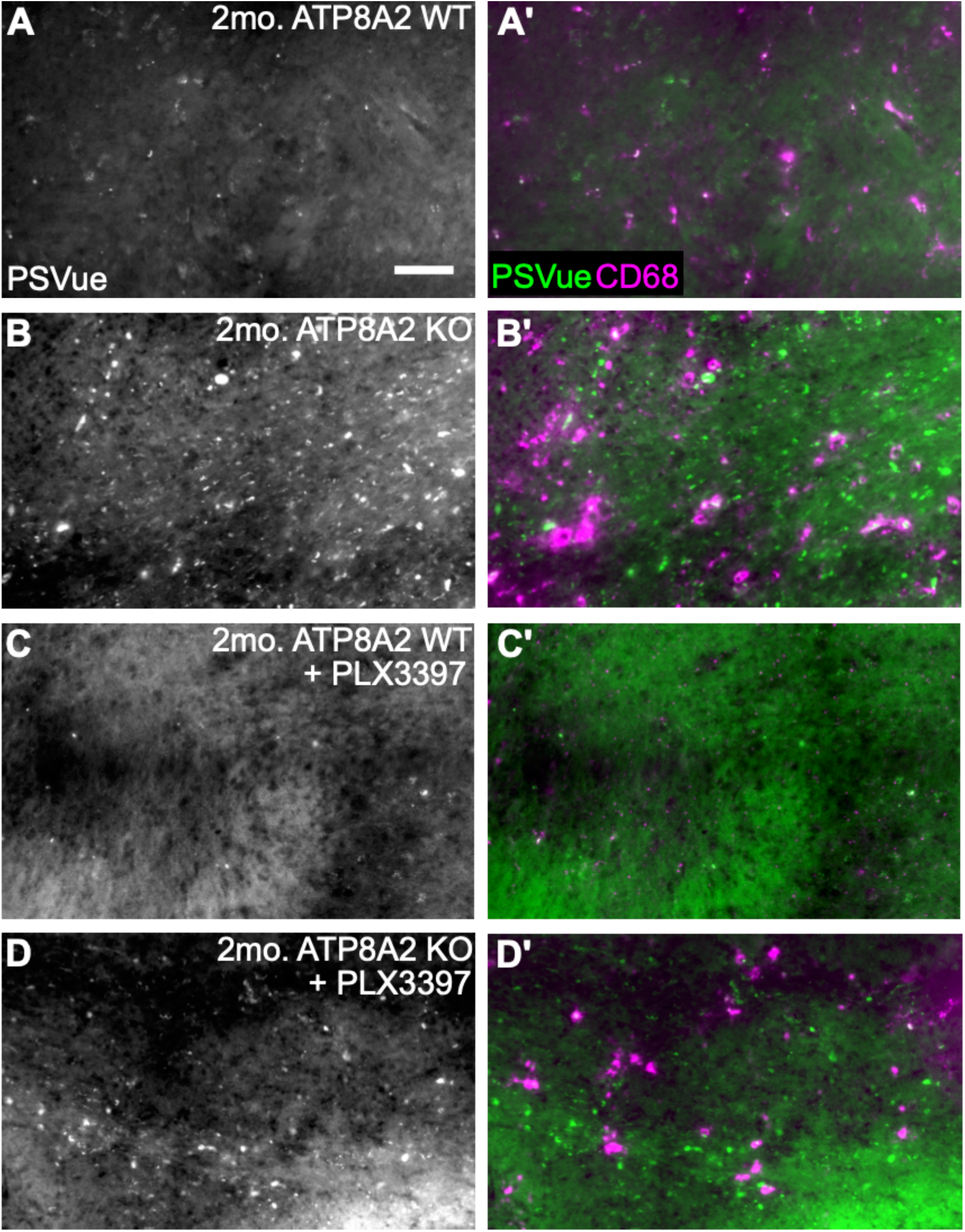
Loss of *Atp8a2* causes externalized phosphatidylserine near the cerebellar peduncles that cannot be rescued by PLX3397. B-E) Brains stained for PSVue-643 (green) and CD68 (magenta) showing an increase in externalized PS in the region of the cerebellar peduncles in 2-month *Atp8a2* knockout mice compared to controls that is not significantly decreased after treatment with 600ppm PLX3397. Scale bar = 50 µm. Quantification in Figure 4E.

**Figure S7:**
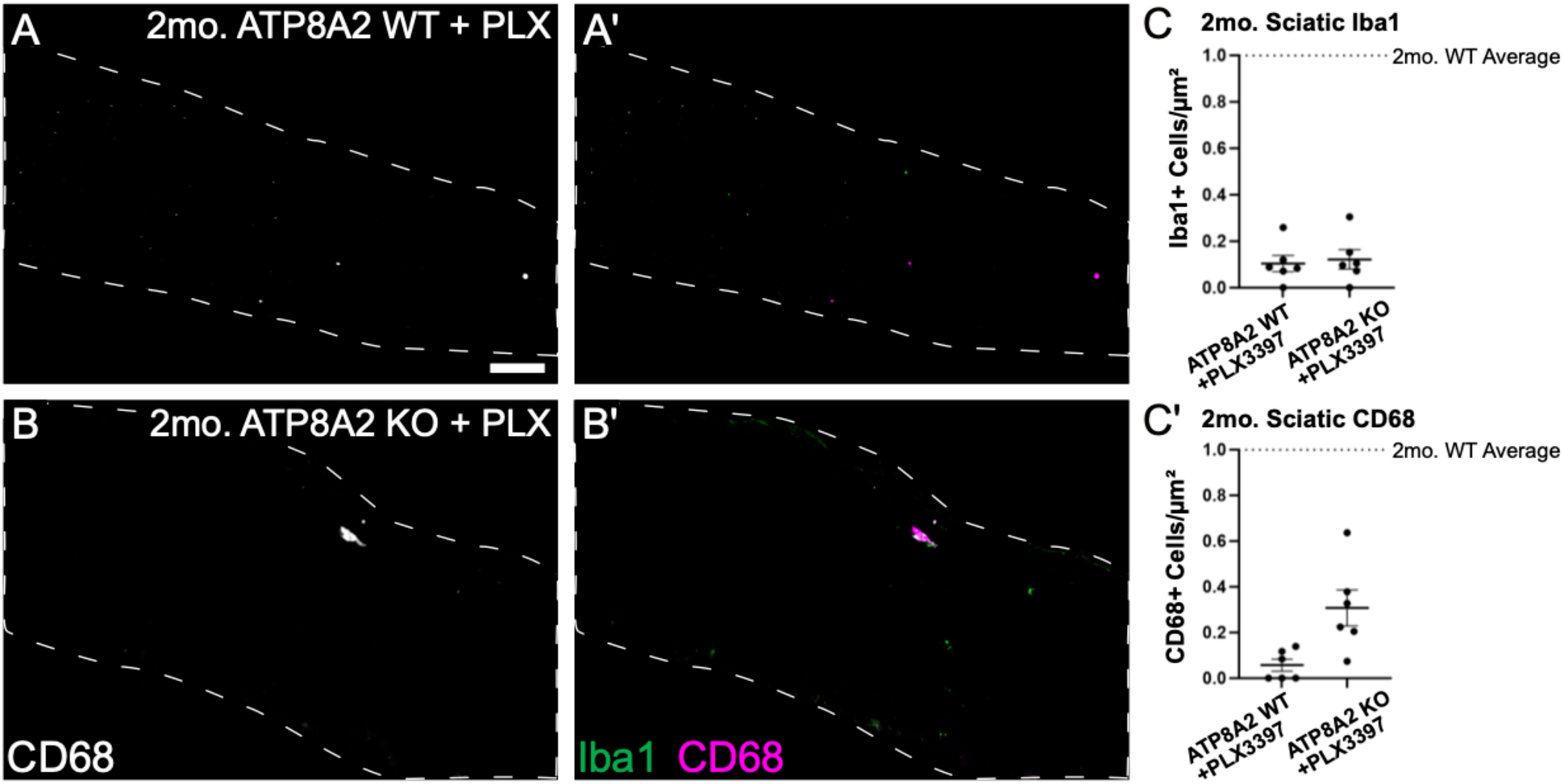
Treatment with PLX3397 depletes peripheral macrophages. A–C) Staining of Sciatic nerves of 2-month mice for Iba1 (pan-macrophages, green) and CD68 (activated macrophages, magenta) shows peripheral macrophage depletion in control (A) and *Atp8a2* knockout mice treated with 600-ppm PLX3397 beginning at Day 30 (B). C) Quantification of Iba1+ cells (C) and CD68+ cells (C’), dotted line represents the mean Iba1+ and CD68+ cells for 2-month control mice in Figure 2D, and D’, respectively. Scale bar = 50 µm.

**Supplemental Table 1:** List of TDP-43-dysregulated genes observed in disparate systems. 23 genes found to be dysregulated in both TDP-43-negative brains of ALS-FTD patients in Ma, et al.^16^ and TDP-43-depleted iPSC-derived cortical-like neurons in Brown, et al.^17^.

| **<u>Gene Symbol</u>** |
| --- |
| *AKT3* |
| *ATP8A2* |
| *CACNA1E* |
| *CADPS* |
| *CAMK2B* |
| *CEP290* |
| *CYFIP2* |
| *ERC2* |
| *EPB41L1* |
| *ICA1* |
| *KALRN* |
| *MADD* |
| *MNAT1* |
| *PLEKHA1* |
| *PRUNE2* |
| *RAP1GAP* |
| *RAPGEF4* |
| *SCN1A* |
| *SHISA9* |
| *STMN2* |
| *STXBP1* |
| *UNC13A* |
| *UQCRC2* |

## References

1. Neumann, M., Sampathu, D.M., Kwong, L.K., Truax, A.C., Micsenyi, M.C., Chou, T.T., Bruce, J., Schuck, T., Grossman, M., Clark, C.M., et al. (2006). Ubiquitinated TDP-43 in Frontotemporal Lobar Degeneration and Amyotrophic Lateral Sclerosis. Science 314, 130–133. 10.1126/science.1134108.

2. Arai, T., Hasegawa, M., Akiyama, H., Ikeda, K., Nonaka, T., Mori, H., Mann, D., Tsuchiya, K., Yoshida, M., Hashizume, Y., et al. (2006). TDP-43 is a component of ubiquitin-positive tau-negative inclusions in frontotemporal lobar degeneration and amyotrophic lateral sclerosis. Biochem. Biophys. Res. Commun. 351, 602–611. 10.1016/j.bbrc.2006.10.093.

3. Igaz, L.M., Kwong, L.K., Lee, E.B., Chen-Plotkin, A., Swanson, E., Unger, T., Malunda, J., Xu, Y., Winton, M.J., Trojanowski, J.Q., et al. (2011). Dysregulation of the ALS-associated gene TDP-43 leads to neuronal death and degeneration in mice. J Clin Invest 121, 726–738. 10.1172/jci44867.

4. Geser, F., Winton, M.J., Kwong, L.K., Xu, Y., Xie, S.X., Igaz, L.M., Garruto, R.M., Perl, D.P., Galasko, D., Lee, V.M.-Y., et al. (2008). Pathological TDP-43 in parkinsonism–dementia complex and amyotrophic lateral sclerosis of Guam. Acta Neuropathol. 115, 133–145. 10.1007/s00401-007-0257-y.

5. Boer, E.M.J. de, Orie, V.K., Williams, T., Baker, M.R., Oliveira, H.M.D., Polvikoski, T., Silsby, M., Menon, P., Bos, M. van den, Halliday, G.M., et al. (2021). TDP-43 proteinopathies: a new wave of neurodegenerative diseases. J Neurology Neurosurg Psychiatry 92, 86–95. 10.1136/jnnp-2020-322983.

6. Meneses, A., Koga, S., O’Leary, J., Dickson, D.W., Bu, G., and Zhao, N. (2021). TDP-43 Pathology in Alzheimer’s Disease. Mol Neurodegener 16, 84. 10.1186/s13024-021-00503-x.

7. Nelson, P.T., Dickson, D.W., Trojanowski, J.Q., Jack, C.R., Boyle, P.A., Arfanakis, K., Rademakers, R., Alafuzoff, I., Attems, J., Brayne, C., et al. (2019). Limbic-predominant age-related TDP-43 encephalopathy (LATE): consensus working group report. Brain 142, 1503–1527. 10.1093/brain/awz099.

8. Jo, M., Lee, S., Jeon, Y.-M., Kim, S., Kwon, Y., and Kim, H.-J. (2020). The role of TDP-43 propagation in neurodegenerative diseases: integrating insights from clinical and experimental studies. Exp. Mol. Med. 52, 1652–1662. 10.1038/s12276-020-00513-7.

9. Mehta, P.R., Brown, A.-L., Ward, M.E., and Fratta, P. (2023). The era of cryptic exons: implications for ALS-FTD. Mol. Neurodegener. 18, 16. 10.1186/s13024-023-00608-5.

10. Ling, J.P., Pletnikova, O., Troncoso, J.C., and Wong, P.C. (2015). TDP-43 repression of nonconserved cryptic exons is compromised in ALS-FTD. Science 349, 650–655. 10.1126/science.aab0983.

11. Klim, J.R., Williams, L.A., Limone, F., Juan, I.G.S., Davis-Dusenbery, B.N., Mordes, D.A., Burberry, A., Steinbaugh, M.J., Gamage, K.K., Kirchner, R., et al. (2019). ALS IMPLICATED PROTEIN TDP-43 SUSTAINS LEVELS OF STMN2 A MEDIATOR OF MOTOR NEURON GROWTH AND REPAIR. Nat Neurosci 22, 167–179. 10.1038/s41593-018-0300-4.

12. Juan, I.G.S., Nash, L.A., Smith, K.S., Leyton-Jaimes, M.F., Qian, M., Klim, J.R., Limone, F., Dorr, A.B., Couto, A., Pintacuda, G., et al. (2022). Loss of mouse Stmn2 function causes motor neuropathy. Neuron 110, 1671–1688.e6. 10.1016/j.neuron.2022.02.011.

13. Krus, K.L., Strickland, A., Yamada, Y., Devault, L., Schmidt, R.E., Bloom, A.J., Milbrandt, J., and DiAntonio, A. (2022). Loss of Stathmin-2, a hallmark of TDP-43-associated ALS, causes motor neuropathy. Cell Reports 39, 111001. 10.1016/j.celrep.2022.111001.

14. Melamed, Z., López-Erauskin, J., Baughn, M.W., Zhang, O., Drenner, K., Sun, Y., Freyermuth, F., McMahon, M.A., Beccari, M.S., Artates, J.W., et al. (2019). Premature polyadenylation-mediated loss of stathmin-2 is a hallmark of TDP-43-dependent neurodegeneration. Nat Neurosci 22, 180–190. 10.1038/s41593-018-0293-z.

15. López-Erauskin, J., Bravo-Hernandez, M., Presa, M., Baughn, M.W., Melamed, Z., Beccari, M.S., Quadros, A.R.A. de A., Arnold-Garcia, O., Zuberi, A., Ling, K., et al. (2024). Stathmin-2 loss leads to neurofilament-dependent axonal collapse driving motor and sensory denervation. Nat. Neurosci. 27, 34–47. 10.1038/s41593-023-01496-0.

16. Ma, X.R., Prudencio, M., Koike, Y., Vatsavayai, S.C., Kim, G., Harbinski, F., Briner, A., Rodriguez, C.M., Guo, C., Akiyama, T., et al. (2022). TDP-43 represses cryptic exon inclusion in the FTD–ALS gene UNC13A. Nature 603, 124–130. 10.1038/s41586-022-04424-7.

17. Brown, A.-L., Wilkins, O.G., Keuss, M.J., Hill, S.E., Zanovello, M., Lee, W.C., Bampton, A., Lee, F.C.Y., Masino, L., Qi, Y.A., et al. (2022). TDP-43 loss and ALS-risk SNPs drive mis-splicing and depletion of UNC13A. Nature 603, 131–137. 10.1038/s41586-022-04436-3.

18. Yang, X., Wang, S., Sheng, Y., Zhang, M., Zou, W., Wu, L., Kang, L., Rizo, J., Zhang, R., Xu, T., et al. (2015). Syntaxin opening by the MUN domain underlies the function of Munc13 in synaptic-vesicle priming. Nat. Struct. Mol. Biol. 22, 547–554. 10.1038/nsmb.3038.

19. Karlsson, M., Zhang, C., Méar, L., Zhong, W., Digre, A., Katona, B., Sjöstedt, E., Butler, L., Odeberg, J., Dusart, P., et al. (2021). A single–cell type transcriptomics map of human tissues. Sci. Adv. 7, eabh2169. 10.1126/sciadv.abh2169.

20. Coleman, J.A., Zhu, X., Djajadi, H.R., Molday, L.L., Smith, R.S., Libby, R.T., John, S.W.M., and Molday, R.S. (2014). Phospholipid flippase ATP8A2 is required for normal visual and auditory function and photoreceptor and spiral ganglion cell survival. J. Cell Sci. 127, 1138–1149. 10.1242/jcs.145052.

21. Zhu, X., Libby, R.T., Vries, W.N. de, Smith, R.S., Wright, D.L., Bronson, R.T., Seburn, K.L., and John, S.W.M. (2012). Mutations in a P-Type ATPase Gene Cause Axonal Degeneration. Plos Genet 8, e1002853. 10.1371/journal.pgen.1002853.

22. Chalat, M., Moleschi, K., and Molday, R.S. (2017). C-terminus of the P4-ATPase ATP8A2 functions in protein folding and regulation of phospholipid flippase activity. Mol. Biol. Cell 28, 452–462. 10.1091/mbc.e16-06-0453.

23. Norris, A.C., Mansueto, A.J., Jimenez, M., Yazlovitskaya, E.M., Jain, B.K., and Graham, T.R. (2024). Flipping the script: Advances in understanding how and why P4-ATPases flip lipid across membranes. Biochim. Biophys. Acta (BBA) - Mol. Cell Res. 1871, 119700. 10.1016/j.bbamcr.2024.119700.

24. Sapar, M.L., Ji, H., Wang, B., Poe, A.R., Dubey, K., Ren, X., Ni, J.-Q., and Han, C. (2018). Phosphatidylserine Externalization Results from and Causes Neurite Degeneration in Drosophila. Cell Reports 24, 2273–2286. 10.1016/j.celrep.2018.07.095.

25. Segawa, K., and Nagata, S. (2015). An Apoptotic ‘Eat Me’ Signal: Phosphatidylserine Exposure. Trends Cell Biol 25, 639–650. 10.1016/j.tcb.2015.08.003.

26. Ravichandran, K.S. (2010). Find-me and eat-me signals in apoptotic cell clearance: progress and conundrums. J Exp Med 207, 1807–1817. 10.1084/jem.20101157.

27. Shacham-Silverberg, V., Shalom, H.S., Goldner, R., Golan-Vaishenker, Y., Gurwicz, N., Gokhman, I., and Yaron, A. (2018). Phosphatidylserine is a marker for axonal debris engulfment but its exposure can be decoupled from degeneration. Cell Death Dis 9, 1116. 10.1038/s41419-018-1155-z.

28. Liu, J., and Wang, F. (2017). Role of Neuroinflammation in Amyotrophic Lateral Sclerosis: Cellular Mechanisms and Therapeutic Implications. Front Immunol 8, 1005. 10.3389/fimmu.2017.01005.

29. Chiot, A., Lobsiger, C.S., and Boillée, S. (2019). New insights on the disease contribution of neuroinflammation in amyotrophic lateral sclerosis. Curr Opin Neurol 32, 764–770. 10.1097/wco.0000000000000729.

30. Lall, D., and Baloh, R.H. (2017). Microglia and C9orf72 in neuroinflammation and ALS and frontotemporal dementia. J. Clin. Investig. 127, 3250–3258. 10.1172/jci90607.

31. Ji, H., Sapar, M.L., Sarkar, A., Wang, B., and Han, C. (2022). Phagocytosis and self-destruction break down dendrites of Drosophila sensory neurons at distinct steps of Wallerian degeneration. Proc National Acad Sci 119, e2111818119. 10.1073/pnas.2111818119.

32. Yang, Y., Sun, K., Liu, W., Zhang, L., Peng, K., Zhang, S., Li, S., Yang, M., Jiang, Z., Lu, F., et al. (2018). Disruption of Tmem30a results in cerebellar ataxia and degeneration of Purkinje cells. Cell Death Dis. 9, 899. 10.1038/s41419-018-0938-6.

33. Ko, K.W., Devault, L., Sasaki, Y., Milbrandt, J., and DiAntonio, A. (2021). Live imaging reveals the cellular events downstream of SARM1 activation. Elife 10, e71148. 10.7554/elife.71148.

34. Dingwall, C.B., Sasaki, Y., Strickland, A., Wu, T., Summers, D.W., Bloom, A.J., DiAntonio, A., and Milbrandt, J. (2025). Suppressing phagocyte activation by overexpressing the phosphatidylserine lipase ABHD12 preserves sarmopathic nerves. iScience 28, 112626. 10.1016/j.isci.2025.112626.

35. Alsahli, S., Alrifai, M.T., Tala, S.A., Mutairi, F.A., and Alfadhel, M. (2018). Further Delineation of the Clinical Phenotype of Cerebellar Ataxia, Mental Retardation, and Disequilibrium Syndrome Type 4. J. Cent. Nerv. Syst. Dis. 10, 1179573518759682. 10.1177/1179573518759682.

36. Guissart, C., Harrison, A.N., Benkirane, M., Oncel, I., Arslan, E.A., Chassevent, A.K.., Baraῆano, K., Larrieu, L., Iascone, M., Tenconi, R., et al. (2020). ATP8A2-related disorders as recessive cerebellar ataxia. J. Neurol. 267, 203–213. 10.1007/s00415-019-09579-4.

37. Cacciagli, P., Haddad, M.-R., Mignon-Ravix, C., El-Waly, B., Moncla, A., Missirian, C., Chabrol, B., and Villard, L. (2010). Disruption of the ATP8A2 gene in a patient with a t(10;13) de novo balanced translocation and a severe neurological phenotype. Eur J Hum Genet 18, 1360–1363. 10.1038/ejhg.2010.126.

38. McMillan, H.J., Telegrafi, A., Singleton, A., Cho, M.T., Lelli, D., Lynn, F.C., Griffin, J., Asamoah, A., Rinne, T., Erasmus, C.E., et al. (2018). Recessive mutations in ATP8A2 cause severe hypotonia, cognitive impairment, hyperkinetic movement disorders and progressive optic atrophy. Orphanet J Rare Dis 13, 86. 10.1186/s13023-018-0825-3.

39. Luse, S.A., Chenard, C., and Finke, E.H. (1967). The wabbler-lethal mouse. An electron microscopic study of the nervous system. Arch. Neurol. 17, 153–161. 10.1001/archneur.1967.00470260043004.

40. Galimberti, D., Bonsi, R., Fenoglio, C., Serpente, M., Cioffi, S.M.G., Fumagalli, G., Arighi, A., Ghezzi, L., Arcaro, M., Mercurio, M., et al. (2015). Inflammatory molecules in Frontotemporal Dementia: Cerebrospinal fluid signature of progranulin mutation carriers. Brain, Behav., Immun. 49, 182–187. 10.1016/j.bbi.2015.05.006.

41. Mackenzie, I.R.A., Baker, M., Pickering-Brown, S., Hsiung, G.-Y.R., Lindholm, C., Dwosh, E., Gass, J., Cannon, A., Rademakers, R., Hutton, M., et al. (2006). The neuropathology of frontotemporal lobar degeneration caused by mutations in the progranulin gene. Brain 129, 3081–3090. 10.1093/brain/awl271.

42. Zhang, J., Velmeshev, D., Hashimoto, K., Huang, Y.-H., Hofmann, J.W., Shi, X., Chen, J., Leidal, A.M., Dishart, J.G., Cahill, M.K., et al. (2020). Neurotoxic microglia promote TDP-43 proteinopathy in progranulin deficiency. Nature 588, 459–465. 10.1038/s41586-020-2709-7.

43. Liu, E.Y., Russ, J., Cali, C.P., Phan, J.M., Amlie-Wolf, A., and Lee, E.B. (2019). Loss of Nuclear TDP-43 Is Associated with Decondensation of LINE Retrotransposons. Cell Rep. 27, 1409–1421.e6. 10.1016/j.celrep.2019.04.003.

44. Kellett, E.A., Bademosi, A.T., and Walker, A.K. (2025). Molecular mechanisms and consequences of TDP-43 phosphorylation in neurodegeneration. Mol. Neurodegener. 20, 53. 10.1186/s13024-025-00839-8.

45. Figley, M.D., and DiAntonio, A. (2020). The SARM1 axon degeneration pathway: control of the NAD+ metabolome regulates axon survival in health and disease. Curr. Opin. Neurobiol. 63, 59–66. 10.1016/j.conb.2020.02.012.

46. MacDonald, K.P.A., Palmer, J.S., Cronau, S., Seppanen, E., Olver, S., Raffelt, N.C., Kuns, R., Pettit, A.R., Clouston, A., Wainwright, B., et al. (2010). An antibody against the colony-stimulating factor 1 receptor depletes the resident subset of monocytes and tissue- and tumor-associated macrophages but does not inhibit inflammation. Blood 116, 3955–3963. 10.1182/blood-2010-02-266296.

47. Pimenova, A.A., Marcora, E., and Goate, A.M. (2017). A Tale of Two Genes: Microglial Apoe and Trem2. Immunity 47, 398–400. 10.1016/j.immuni.2017.08.015.

48. Makarava, N., Safadi, T., Bocharova, O., Mychko, O., Pandit, N.P., Molesworth, K., Eyo, U.B., and Baskakov, I.V. (2025). Knockout of P2Y12 receptor facilitates neuronal envelopment by reactive microglia and accelerates prion disease. J. Neuroinflammation 22, 210. 10.1186/s12974-025-03542-z.

49. Yang, S., Wang, J., Cao, Y., Zhang, Y., Sun, Z., Wan, P., Pi, M., Xiong, Q., Shu, X., Wang, X., et al. (2025). Clec7a Signaling in Microglia Promotes Synapse Loss Associated with Tauopathy. Int. J. Mol. Sci. 26, 2888. 10.3390/ijms26072888.

50. Kebschull, J.M., Richman, E.B., Ringach, N., Friedmann, D., Albarran, E., Kolluru, S.S., Jones, R.C., Allen, W.E., Wang, Y., Cho, S.W., et al. (2020). Cerebellar nuclei evolved by repeatedly duplicating a conserved cell-type set. Science 370. 10.1126/science.abd5059.

51. Elmore, M.R.P., Najafi, A.R., Koike, M.A., Dagher, N.N., Spangenberg, E.E., Rice, R.A., Kitazawa, M., Matusow, B., Nguyen, H., West, B.L., et al. (2014). Colony-Stimulating Factor 1 Receptor Signaling Is Necessary for Microglia Viability, Unmasking a Microglia Progenitor Cell in the Adult Brain. Neuron 82, 380–397. 10.1016/j.neuron.2014.02.040.

52. Hohsfield, L.A., Najafi, A.R., Ghorbanian, Y., Soni, N., Crapser, J., Velez, D.X.F., Jiang, S., Royer, S.E., Kim, S.J., Henningfield, C.M., et al. (2021). Subventricular zone/white matter microglia reconstitute the empty adult microglial niche in a dynamic wave. eLife 10, e66738. 10.7554/elife.66738.

53. Groh, J., Klein, D., Berve, K., West, B.L., and Martini, R. (2019). Targeting microglia attenuates neuroinflammation-related neural damage in mice carrying human PLP1 mutations. Glia 67, 277–290. 10.1002/glia.23539.

54. Tollervey, J.R., Curk, T., Rogelj, B., Briese, M., Cereda, M., Kayikci, M., König, J., Hortobágyi, T., Nishimura, A.L., Župunski, V., et al. (2011). Characterizing the RNA targets and position-dependent splicing regulation by TDP-43. Nat. Neurosci. 14, 452–458. 10.1038/nn.2778.

55. Polymenidou, M., Lagier-Tourenne, C., Hutt, K.R., Huelga, S.C., Moran, J., Liang, T.Y., Ling, S.-C., Sun, E., Wancewicz, E., Mazur, C., et al. (2011). Long pre-mRNA depletion and RNA missplicing contribute to neuronal vulnerability from loss of TDP-43. Nat. Neurosci. 14, 459–468. 10.1038/nn.2779.

56. Joseph, B.J., Marshall, K.A., Harley, P., Mann, J.R., Alessandrini, F., Vanoye, C.G., Chi, W., Prudencio, M., Simkin, D., Kao, T.-T., et al. (2025). TDP-43-dependent mis-splicing of KCNQ2 triggers intrinsic neuronal hyperexcitability in ALS/FTD. Nat. Neurosci., 1–17. 10.1038/s41593-025-02096-w.

57. Beccari, M.S., Arnold-Garcia, O., Baughn, M.W., Artates, J.W., McAlonis-Downes, M., Lim, J., Leyva-Cázares, D.F., Rubio-Lara, H.I., Ramirez-Rodriguez, A., Bernal-Buenrostro, C.N., et al. (2025). Stathmin-2 enhances motor axon regeneration after injury independent of its binding to tubulin. Proc. Natl. Acad. Sci. 122, e2502294122. 10.1073/pnas.2502294122.

58. Li, Y., Tian, Y., Pei, X., Zheng, P., Miao, L., Li, L., Luo, C., Zhang, P., Jiang, B., Teng, J., et al. (2023). SCG10 is required for peripheral axon maintenance and regeneration in mice. J. Cell Sci. 136. 10.1242/jcs.260490.

59. Martín-Hernández, E., Rodríguez-García, M.E., Camacho, A., Matilla-Dueñas, A., García-Silva, M.T., Quijada-Fraile, P., Corral-Juan, M., Tejada-Palacios, P., Heras, R.S. de L., Arenas, J., et al. (2016). New ATP8A2 gene mutations associated with a novel syndrome: encephalopathy, intellectual disability, severe hypotonia, chorea and optic atrophy. neurogenetics 17, 259–263. 10.1007/s10048-016-0496-y.

60. Mendez, J.S., Cohen, A.L., Eckenstein, M., Jensen, R.L., Burt, L.M., Salzman, K.L., Chamberlain, M., Hsu, H.H., Hutchinson, M., Iwamoto, F., et al. (2024). Phase 1b/2 study of orally administered pexidartinib in combination with radiation therapy and temozolomide in patients with newly diagnosed glioblastoma. Neuro-Oncol. Adv. 6, vdae202. 10.1093/noajnl/vdae202.

61. Butowski, N., Colman, H., Groot, J.F.D., Omuro, A.M., Nayak, L., Wen, P.Y., Cloughesy, T.F., Marimuthu, A., Haidar, S., Perry, A., et al. (2015). Orally administered colony stimulating factor 1 receptor inhibitor PLX3397 in recurrent glioblastoma: an Ivy Foundation Early Phase Clinical Trials Consortium phase II study. Neuro-Oncol. 18, 557–564. 10.1093/neuonc/nov245.

62. Terryn, J., Verfaillie, C.M., and Damme, P.V. (2021). Tweaking Progranulin Expression: Therapeutic Avenues and Opportunities. Front. Mol. Neurosci. 14, 713031. 10.3389/fnmol.2021.713031.

63. O’Rourke, J.G., Bogdanik, L., Yáñez, A., Lall, D., Wolf, A.J., Muhammad, A.K.M.G., Ho, R., Carmona, S., Vit, J.P., Zarrow, J., et al. (2016). C9orf72 is required for proper macrophage and microglial function in mice. Science 351, 1324–1329. 10.1126/science.aaf1064.

64. Oeckl, P., Weydt, P., Steinacker, P., Anderl-Straub, S., Nordin, F., Volk, A.E., Diehl-Schmid, J., Andersen, P.M., Kornhuber, J., Danek, A., et al. (2019). Different neuroinflammatory profile in amyotrophic lateral sclerosis and frontotemporal dementia is linked to the clinical phase. J. Neurol., Neurosurg. Psychiatry 90, 4. 10.1136/jnnp-2018-318868.

65. Martens, L.H., Zhang, J., Barmada, S.J., Zhou, P., Kamiya, S., Sun, B., Min, S.-W., Gan, L., Finkbeiner, S., Huang, E.J., et al. (2012). Progranulin deficiency promotes neuroinflammation and neuron loss following toxin-induced injury. J. Clin. Investig. 122, 3955–3959. 10.1172/jci63113.

66. Beel, S., Moisse, M., Damme, M., Muynck, L.D., Robberecht, W., Bosch, L.V.D., Saftig, P., and Damme, P.V. (2017). Progranulin functions as a cathepsin D chaperone to stimulate axonal outgrowth in vivo. Hum. Mol. Genet. 26, 2850–2863. 10.1093/hmg/ddx162.

67. Augustin, I., Rosenmund, C., Südhof, T.C., and Brose, N. (1999). Munc13-1 is essential for fusion competence of glutamatergic synaptic vesicles. Nature 400, 457–461. 10.1038/22768.

68. Varoqueaux, F., Sigler, A., Rhee, J.-S., Brose, N., Enk, C., Reim, K., and Rosenmund, C. (2002). Total arrest of spontaneous and evoked synaptic transmission but normal synaptogenesis in the absence of Munc13-mediated vesicle priming. Proc. Natl. Acad. Sci. 99, 9037–9042. 10.1073/pnas.122623799.

69. Daele, S.H.V., Masrori, P., Damme, P.V., and Bosch, L.V.D. (2024). The sense of antisense therapies in ALS. Trends Mol. Med. 30, 252–262. 10.1016/j.molmed.2023.12.003.

70. Liu, E., Karpf, L., and Bohl, D. (2021). Neuroinflammation in Amyotrophic Lateral Sclerosis and Frontotemporal Dementia and the Interest of Induced Pluripotent Stem Cells to Study Immune Cells Interactions With Neurons. Front. Mol. Neurosci. 14, 767041. 10.3389/fnmol.2021.767041.

71. Chiot, A., Zaïdi, S., Iltis, C., Ribon, M., Berriat, F., Schiaffino, L., Jolly, A., Grange, P. de la, Mallat, M., Bohl, D., et al. (2020). Modifying macrophages at the periphery has the capacity to change microglial reactivity and to extend ALS survival. Nat Neurosci 23, 1339–1351. 10.1038/s41593-020-00718-z.

72. Gilley, J., Ribchester, R.R., and Coleman, M.P. (2017). Sarm1 Deletion, but Not Wld S, Confers Lifelong Rescue in a Mouse Model of Severe Axonopathy. Cell Rep. 21, 10–16. 10.1016/j.celrep.2017.09.027.

73. Sasaki, Y., Nakagawa, T., Mao, X., DiAntonio, A., and Milbrandt, J. (2016). NMNAT1 inhibits axon degeneration via blockade of SARM1-mediated NAD+ depletion. eLife 5, e19749. 10.7554/elife.19749.

74. Beirowski, B., Babetto, E., Gilley, J., Mazzola, F., Conforti, L., Janeckova, L., Magni, G., Ribchester, R.R., and Coleman, M.P. (2009). Non-Nuclear WldS Determines Its Neuroprotective Efficacy for Axons and Synapses In Vivo. J. Neurosci. 29, 653–668. 10.1523/jneurosci.3814-08.2009.

75. Mack, T.G.A., Reiner, M., Beirowski, B., Mi, W., Emanuelli, M., Wagner, D., Thomson, D., Gillingwater, T., Court, F., Conforti, L., et al. (2001). Wallerian degeneration of injured axons and synapses is delayed by a Ube4b/Nmnat chimeric gene. Nat. Neurosci. 4, 1199–1206. 10.1038/nn770.

76. Rojo, R., Raper, A., Ozdemir, D.D., Lefevre, L., Grabert, K., Wollscheid-Lengeling, E., Bradford, B., Caruso, M., Gazova, I., Sánchez, A., et al. (2019). Deletion of a Csf1r enhancer selectively impacts CSF1R expression and development of tissue macrophage populations. Nat. Commun. 10, 3215. 10.1038/s41467-019-11053-8.

77. Prudencio, M., Humphrey, J., Pickles, S., Brown, A.-L., Hill, S.E., Kachergus, J.M., Shi, J., Heckman, M.G., Spiegel, M.R., Cook, C., et al. (2020). Truncated stathmin-2 is a marker of TDP-43 pathology in frontotemporal dementia. J. Clin. Investig. 130, 6080–6092. 10.1172/jci139741.

78. Schindelin, J., Arganda-Carreras, I., Frise, E., Kaynig, V., Longair, M., Pietzsch, T., Preibisch, S., Rueden, C., Saalfeld, S., Schmid, B., et al. (2012). Fiji: an open-source platform for biological-image analysis. Nat Methods 9, 676–682. 10.1038/nmeth.2019.

79. Li, Y.I., Knowles, D.A., Humphrey, J., Barbeira, A.N., Dickinson, S.P., Im, H.K., and Pritchard, J.K. (2018). Annotation-free quantification of RNA splicing using LeafCutter. Nat. Genet. 50, 151–158. 10.1038/s41588-017-0004-9.

80. Kaiser, T., Allen, H.M., Kwon, O., Barak, B., Wang, J., He, Z., Jiang, M., and Feng, G. (2021). MyelTracer: A Semi-Automated Software for Myelin g-Ratio Quantification. eNeuro 8, ENEURO.0558-20.2021. 10.1523/eneuro.0558-20.2021.

81. Szretter, K.J., Samuel, M.A., Gilfillan, S., Fuchs, A., Colonna, M., and Diamond, M.S. (2009). The Immune Adaptor Molecule SARM Modulates Tumor Necrosis Factor Alpha Production and Microglia Activation in the Brainstem and Restricts West Nile Virus Pathogenesis. J. Virol. 83, 9329–9338. 10.1128/jvi.00836-09.

82. Cai, W., Haddad, M., Haddad, R., Kesten, I., Hoffman, T., Laan, R., Westfall, S., Defaye, M., Abdullah, N.S., Wong, C., et al. (2025). The gut microbiota promotes pain in fibromyalgia. Neuron 113, 2161–2175.e13. 10.1016/j.neuron.2025.03.032.

83. Lee, S., Shi, X.Q., Fan, A., West, B., and Zhang, J. (2018). Targeting macrophage and microglia activation with colony stimulating factor 1 receptor inhibitor is an effective strategy to treat injury-triggered neuropathic pain. Mol. Pain 14, 1744806918764979. 10.1177/1744806918764979.

84. Luong, T.N., Carlisle, H.J., Southwell, A., and Patterson, P.H. (2011). Assessment of Motor Balance and Coordination in Mice using the Balance Beam. J. Vis. Exp. 10.3791/2376.

85. Zeng, Y., Sianto, O., Lovchykova, A., Liu, C., Akiyama, T., Petrucelli, L., and Gitler, A.D. (2025). Nonsense-mediated decay masks cryptic splicing events caused by TDP-43 loss. bioRxiv, 2025.07.09.664014. 10.1101/2025.07.09.664014.

